# Spatially heterogeneous selection and inter-varietal differentiation maintain population structure and local adaptation in a widespread conifer

**DOI:** 10.1101/2024.04.03.587947

**Authors:** Pablo Peláez, Gustavo P. Lorenzana, Kailey Baesen, Jose Ruben Montes, Amanda R. De La Torre

## Abstract

Douglas-fir (*Pseudotsuga menziesii*) plays a critical role in the ecology and economy of Western North America. This conifer species comprises two distinct varieties: the coastal variety (var. *menziesii*) along the Pacific coast, and the interior variety (var. *glauca*) spanning the Rocky Mountains into Mexico, with instances of inter-varietal hybridization in Washington and British Columbia. Recent investigations have focused on assessing environmental pressures shaping Douglas-fir’s genomic variation for a better understanding of its evolutionary and adaptive responses. Here, we characterize range-wide population structure, estimate inter-varietal hybridization levels, identify candidate loci for climate adaptation, and forecast shifts in species and variety distribution under future climates.

Using a custom SNP-array, we genotyped 540 trees revealing four distinct clusters with asymmetric admixture patterns in the hybridization zone. Higher genetic diversity observed in coastal and hybrid populations contrasts with lower diversity in inland populations of the southern Rockies and Mexico, exhibiting a significant isolation by distance pattern, with less marked but still significant isolation by environment. For both varieties, we identified candidate loci associated with local adaptation, with hundreds of genes linked to processes such as stimulus response, reactions to chemical compounds, and metabolic functions. Ecological niche modeling showed contrasting potential distribution shifts among varieties in the coming decades.

Overall, our findings provide crucial insights into the population structure and adaptive potential of Douglas-fir, with the coastal variety being the most likely to preserve its evolutionary path throughout the present century, which carry implications for the conservation and management of this species across their range.

## Background

Intraspecific gene flow can generate novel genetic pools and promote adaptive evolution in differentiated populations. Intraspecific variation and hybridization in natural populations with dissimilar evolutionary histories are widely acknowledged as crucial factors for different patterns of local adaptation, and understanding these evolutionary processes has become more relevant due to climate change [1]. Forest trees have been proposed to be susceptible to rapid climate change due to their long generation times. However, trees also present high levels of standing genetic diversity due to large populations sizes, extensive levels of gene flow and wide distributions across different environments, which makes them ideal systems to detect adaptive signals and to study the genetic basis of adaptation [2]. In the past, efforts to dissect the genetic basis of adaptation relied on population genetic analyses that focused mainly on the genetic differentiation of the populations without considering environmental heterogeneity among them [3]. Recent approaches in landscape genomics integrate multiple genetic, spatial, and environmental data to shed light on the genetic variants underlying adaptation to different climate scenarios.

Douglas-fir constitutes one of the world’s most important timber trees. It grows under a wide range of climatic conditions and has been part of the landscape of western North America since the Pleistocene. With warming environmental conditions, research on Douglas-fir has focused on testing the ability of the species to grow while withstanding changes in temperature (heat stress) and low soil water content (drought stress) [4,5,6,7,8]. Two varieties with adaptive variation are formally recognized: the coastal variety along the Pacific coast (*P. menziesii* var. *menziesii*), and the interior variety across the Rocky Mountains (*P. menziesii* var. *glauca*).

Populations of Mexican Douglas fir are small, extremely fragmented, and non-continuous in distribution, except in a larger area in Chihuahua [9]. Anthropogenic pressure, as well as the high rate of deforestation, threaten several Douglas fir populations and other conifers, reducing their size and variability within populations [10,11]. Consequently, the Mexican government has listed the Mexican Douglas fir as rare and subject to special protection. Compared to populations from northern Mexico, populations from central Mexico are genetically, morphologically, and phenologically more distinct from populations from the United States [12,13,14,15]. Particularly in Mexico, there is high genetic diversity among Douglas fir populations [14,15]. Despite its genetic and ecological importance, there are no conservation efforts for the Mexican Douglas fir in Mexico. Some studies have suggested considering patterns of genetic diversity and population history, *in-situ* and *ex-situ* conservation activities, and germplasm collected for assisted gene flow and migration to reduce inbreeding [16,17].

Fossil records exist for Douglas-fir from the early Miocene to the late Holocene [15]. Fossils from the Miocene and Pliocene were located along the west coast (from British Columbia to California) and in the Columbia Plateau and Great Basin. Pleistocene fossils were detected in the Rocky Mountains and in the west coast; however, no fossil records were found in the Columbia Plateau and the western Great Basin [15]. This change in the distribution of fossil records correlates with the Pliocene orogeny of the Sierra Nevada and Cascade Mountain ranges, which is thought to be the cause of the vicariant separation of the species into interior and coastal varieties [15]. In addition, fossil records suggest that subsequent differentiation within varieties may have occurred due to climate change refugia during the Pleistocene glaciation [15,18]. Posterior events of secondary contact in British Columbia and the Washington Cascades led to the formation of two inter-varietal hybrid zones in these geographic locations. The extent and distribution of hybridization in the species is unknown.

Demographic, phylogenetic and population molecular studies have shown differentiation between the varieties and populations of Douglas-fir [15,19,20,21,22]. Genetic differentiation between the two varieties was reported using allozymes more than three decades ago [20]. Chloroplast and mitochondrial markers revealed Pleistocene divergence of the Mexican populations, which resulted in the classification of the interior variety into two different lineages: the Rocky Mountain and Mexican lineages [15,19]. Population structure analyses using nuclear microsatellite markers identified refugial populations with intra-varietal and inter-varietal gene flow caused by hybridization and introgression [18,22].

Studies that assess the relation between genetic diversity and environmental responses in Douglas-fir have been scarce. Recently, based on genome-wide sequencing approaches, genes associated with local adaptation have been identified [23,24,25,7]. A comparative transcriptome analysis of Douglas-fir trees from two provenances, from a coastal and interior habitat, with contrasting natural environments was carried out to evaluate the effect of abiotic environmental factors on gene expression responses [24]. Targeted sequence capture and mixed effect models were used to detect high differentiation of drought tolerance genes between interior and coastal trees grown in experimental conditions [25]. In coastal populations, single nucleotide polymorphisms have been associated with cold-hardiness and phenology related traits [23,7]. In this study, we genotyped trees of the two varieties to characterize the population structure across the natural, species-range distribution of Douglas fir in North America; identify the evolutionary processes maintaining genetic variation and population structure; assess the levels of inter-varietal hybridization; identify candidate loci for climate adaptation; and predict the impacts of climate change in the future distribution of the species.

## Materials and Methods

### Plant material and DNA extraction

Seeds and needle tissue were collected from 577 open-pollinated trees across the Douglas-fir natural distribution range, from Mexico to British Columbia, Canada (Table 1). Prior to extraction, seeds were soaked in a solution of 70% water and 30% of 3% hydrogen peroxide for 12 hrs. Ten haploid megagametophytes for each family were pooled to infer the maternal genotype. DNA from megagametophytes was extracted using the Qiagen DNeasy mini-prep Plant kit and an Eppendorf automated pipetting workstation. When needle tissue was available, DNA was extracted using a modified CTAB method [26] or the MPBio (MPBiomedicals LLC, Ohio, USA) RapidPure DNA Plant kit. The CTAB method was modified to include a wash step of homogenized tissue with CTAB buffer prior to 65°C incubation. DNA quality and concentration was evaluated with a NanoDrop Spectrophotometer, a Qubit 4.0 Fluorometer and agarose gels using Invitrogen’s E-Gel Power Snap system.

**Table 1.** Geographic location and mean environmental variables of individuals included in this study.

| Group | Latitude | Longitude | MAT (°C) | MAP (mm) | Sample size |
| --- | --- | --- | --- | --- | --- |
| Coastal | 45.8 | -123.0 | 8.93 | 1861.2 | 325 |
| Hybrid | 51.4 | -120.2 | 3.11 | 752.2 | 27 |
| Interior north | 47.9 | -11.87 | 4.02 | 793.5 | 61 |
| Interior south | 35.5 | -108.3 | 6.88 | 584.1 | 127 |

MAT, mean annual temperature; MAP, mean annual precipitation.

### SNP genotyping and filtering

Samples were genotyped using a custom-designed gene-based Illumina Infinium SNP array containing 16,146 Douglas-fir single nucleotide polymorphisms (SNPs) at the University of California-Davis Genome Center. This array was designed to represent genome-wide variation in the species by using 10X whole-genome re-sequencing data from individuals across the species’ geographic range as input for array construction [7,8]. Genotypes were called using Illumina’s Genome Studio Genotyping Module v 2.0.5 (Illumina, 2016). Filtering criteria included a SNP call frequency > 0.85, individual call rate > 0.85, non-monomorphic and a minor allele frequency > 0.01.

### Population structure and hybridization analyses

Principal component analysis (PCA) was carried out to infer population structure using the gdsfmt v.1.34 and SNPRelate v.1.32 packages [27]. Population structure was also inferred by a Discriminant Analysis of Principal Components using the find.clusters function of the adegenet (v.2.1.10) package [28]. The best number of genetic clusters was determined using the Bayesian Information Criterion (BIC). To estimate ancestral differences between varieties and populations, ADMIXTURE software v.1.3.0 [29] was employed using 10 independent runs for k-values ranging from 2 to 10. Admixture proportions obtained for K=2 and the two K-values with the lowest averaged cross-validation error were chosen. CLUMPAK v.1.1 was used to produce bar plots [30]. Pie chart maps of individual admixture proportions were constructed with QGIS and the option around the point for better visualization. Posterior probabilities of hybrid categories were calculated with NewHybrids v.2.0 [31]. One hundred randomly selected SNPs were used to estimate posterior probabilities with 50,000 sweeps for 3 replicates.

### Genetic diversity analyses

Population inbreeding coefficients were obtained from a kinship matrix with the popkin package v.1.3.23 [32]. Geographic positions of populations on the map were estimated from the centroid of all individuals in each population. The natural distribution map of the two varieties was based on the Atlas of United States trees [33]. The VCFtools program v.0.1.16 was used to determine the heterozygosity and pairwise fixation index (Fst) values [34]. Heterozygosity values were calculated per individual with the observed number of homozygotes [O(HOM)] and the number of non-missing genotypes [N(NM)] using the formula: [N(NM) – O(HOM)]/N(NM). Violin plots were created to visualize the results using the ggplot2 v. 3.4.2 R package [35].

### Isolation by distance and environment analyses

To evaluate the association between genetic and geographic distances, a Mantel test was performed using the gl.ibd function implemented in the dartR R package v. 1.0.2 with 999 permutations [36]. Isolation by environment and association between geographic and environmental distances analyses were performed with vegan (v. 2.6-4) and geosphere (v. 1.5-18) R packages. The Spearman correlation method, 9999 permutations, and SNPs without missing data were used for Mantel tests. Climatic data for the environmental variables (BIO1 and BIO12) were obtained from the WorldClim database (www.worldclim.org) with a resolution of 2.5 arcsec using the raster and sp R packages and the coordinates of the trees collected.

### Detection of loci under selection

Identification of SNPs under selection was carried out with BayeScan (v. 2.0) and pcadapt (v. 4.3.3) packages [37,38]. The parameters used for outliers’ detection with BayeScan were the following: 20 pilot runs of 5000 iterations, burn-in length of 50,000, thinning interval size of 100 and prior odds ratio of 100. SNPs were considered outliers based on a false discovery rate q-value threshold of 0.05. Pcadapt was used to detect signatures of selection using two principal components. A q-value threshold of 0.1 was used to choose outliers. SNPs were functionally annotated using KOBAS-i and the annotation file of the Douglas-fir genome v. 1.0 [39]. KOBAS-i was used with default parameters for Gene Ontology (GO) and KEGG pathways enrichment analyses. Climate data of 25 environmental variables were extracted from the ClimateNA application [40]. All climate data are based on annual averages for the years 1962–1990. The variables used in the analysis are related to monthly, seasonal, and annual temperature and precipitation measurements (Additional file 1). To identify SNPs involved in local adaptation to climate, the Bayenv2 program was used [41]. A covariance matrix was obtained from the average of 5 independent covariance matrices generated with different seed numbers and 100,000 iterations. Five independent runs were performed for the populations of Douglas-fir and 25 climate variables with different seed sizes and 500,000 iterations. Bayes factors (BF) were averaged over the independent runs. A BF value of 10 was considered as threshold for outlier loci.

### Ecological niche modeling (ENM)

The ecological niche of Douglas-fir was modeled with the Wallace2 interactive web app [42], which incorporates the Maxent algorithm for estimating potential distribution using presence-only data [43,44]. Additionally, ecological niche shifts were forecasted (i.e., potential distribution gains and losses in response to climate change) throughout the present century (years 2050 and 2070). Douglas-fir’s presence data was derived from the 540 geo-referenced trees retained after the SNP filtering procedure. From those, 236 duplicated coordinates were further removed. The geographic database was complemented with Douglas-fir site records (n=44) reported by Wei et al. (2011), which allowed to fill some geographic gaps, especially for Mexican populations, increasing the overall presence data across the species range. The resulting 348 non-duplicated locations were then subdivided into three major regions (ancestral populations) according to PCA and admixture results: coastal (n=250), interior-north (n=41), interior-south (n=48), and a hybrid zone (n=9). Coastal region included northern California to British Columbia; interior-north encompassed Idaho and Wyoming up north to British Columbia, and interior-south comprised high elevations from Utah and Colorado to southern Mexico. Hybrid zone is located in British Columbia and Washington.

Nineteen bioclimatic variables derived from present-day averaged temperature and precipitation data available at WorldClim [45] were used as raster layers with a resolution of 2.5 arc minutes (≈ 5 km). Occurrence data was further processed to reduce sampling bias using a spatial thinning technique, where the minimum distance between occurrence locations (i.e., nearest neighbor distance) was 10 km, resulting in a thinned dataset of 263 localities. From those points, the spatial extent for niche model building and evaluation was determined, using a bounding box with a buffer distance of 1 geographic degree and 10,000 sample background points. The environmental space occupied by each of the regional Douglas-fir populations was characterized as an approximation to their Hutchinsonian niche, defined as “the n-dimensional hypervolume where a species can persist and reproduce in a mathematical space defined by non-depletable environmental gradients” [42,46]. Wallace2 reduces the dimensionality of the predicted niche using three modules: 1) “environmental ordination”, which is basically a Principal Component Analysis (PCA), 2) “occurrence density grid”, depicting the portion of the environmental space that is more densely occupied by the species, given the availability of environmental conditions present within the background extent, and 3) “niche overlap”, quantified as an overall index (overlap D) [47], ranging from 0 to 1, where 0 represents a null overlap and 1 is a complete ecological overlap between populations.

A spatial partition of the occurrence data for training and testing the predictive models, was performed using the “checkerboard 1” option, with k=2, and aggregation factor 2. Then, niche models were built using the Maxent module. Maxent is a machine learning algorithm that models the potential distribution of a species in response to a set of environmental conditions (e.g., climate), which are constrained to be relatively uniform across the geographic space encompassed by the input locations [43,48]. Increasingly complex models were obtained using linear, quadratic, hinge, and product feature classes, and their predictive performance was evaluated using a combination of metrics included in Wallace2 such as the Area Under the Curve (AUC), omission rate, Continuous Boyce Index (CBI), and Akaike Information Criterion (AIC). The best models per species and per variety were transferred to geographic space within the range of Douglas-fir in North America, and as well to future climate scenarios over the next 50 years.

## Results

### Data collection and genotyping

Assessments of genetic diversity, population structure and natural hybridization of Douglas-fir were based on the genotypes of 577 trees, grouped in 37 populations across the coastal, interior, Mexican and inter-variety hybrid zones in the species’ natural geographic range (Figure 1; Additional file 2). After filtering steps (SNP and sample call frequency rates and minor allele frequency), 540 trees and 11,320 SNPs representing 70.1% of the SNPs in the array were kept for further analyses. The total genotype rate was 96%.

**Figure 1.**
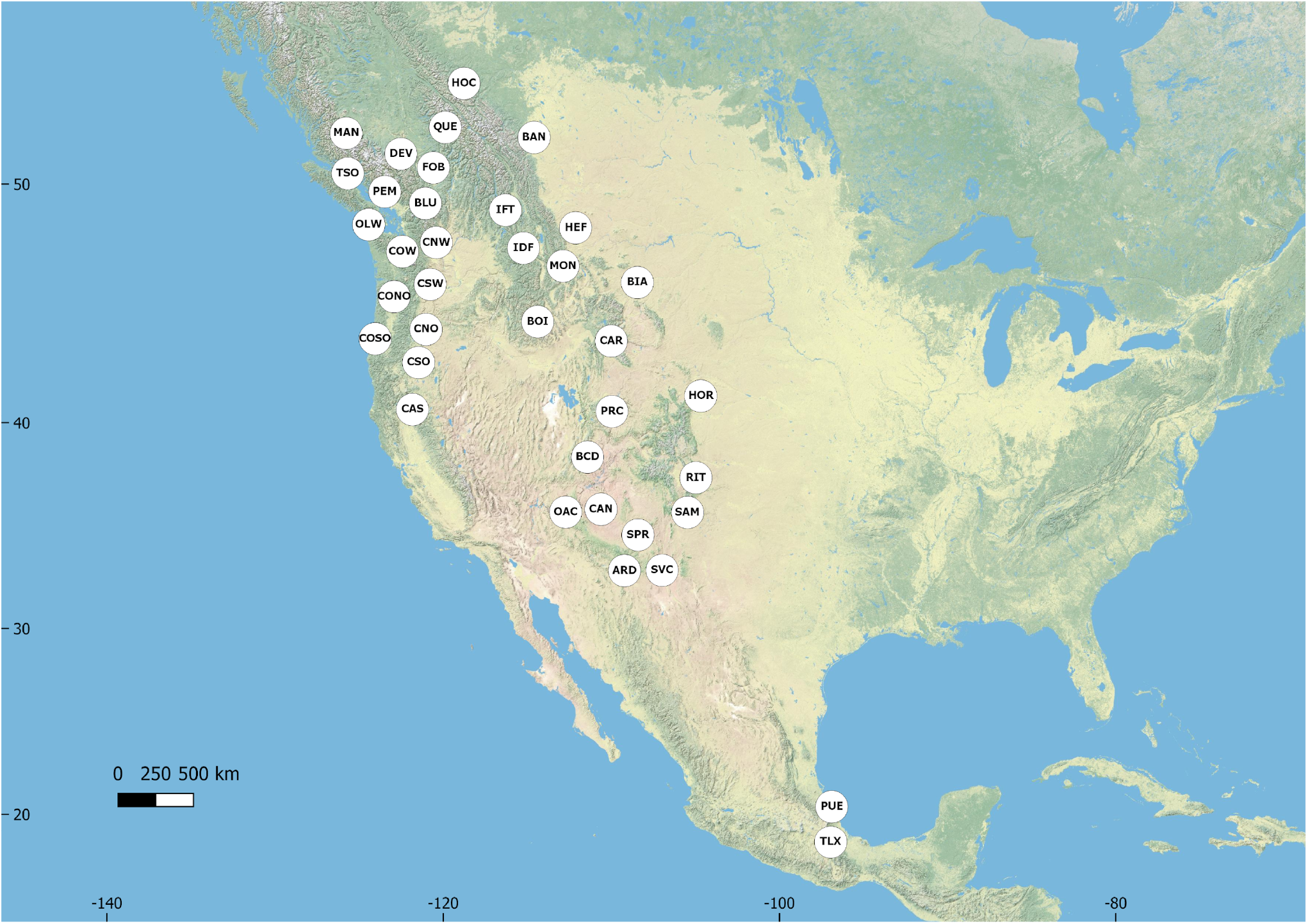
Douglas-fir sampling localities, from Mexico to British Columbia. Circles represent each of the 37 sampled localities.

### Population structure and hybridization

Due to the broad geographic distribution of the two recognized varieties and the presence of contact zones, this study began by determining the population structure of the dataset. Principal component analysis with the SNPRelate package [27] distinguished four genetic clusters, dividing individuals, and populations from coastal, hybrids, interior-north, and interior-south groups/varieties (Figure 2A). Principal component 1, which accounted for 16.7% of the variation in the dataset, separated coastal populations from interior and hybrids, suggesting coastal and interior-south are the most genetically differentiated clusters. Individuals from the Mexican populations clustered together with populations from the interior-south. Principal component 2, which accounted for 4.15% of the variation in the dataset, separated interior-north and hybrids from most interior-south and coastal individuals (Figure 2B). Discriminant analysis of principal components (DAPC) results coincided with PCA results in the optimal number of clusters (Figure 2C). However, the interior and hybrid populations were clustered differently than in the PCA results. Individuals from interior-north and hybrids were clustered together, while interior-south individuals were separated into two clusters that distinguished trees from Arizona, New Mexico, Puebla (Mexico), and Tlaxcala (Mexico), from those in Utah and Colorado (Figure 2D).

**Figure 2.**
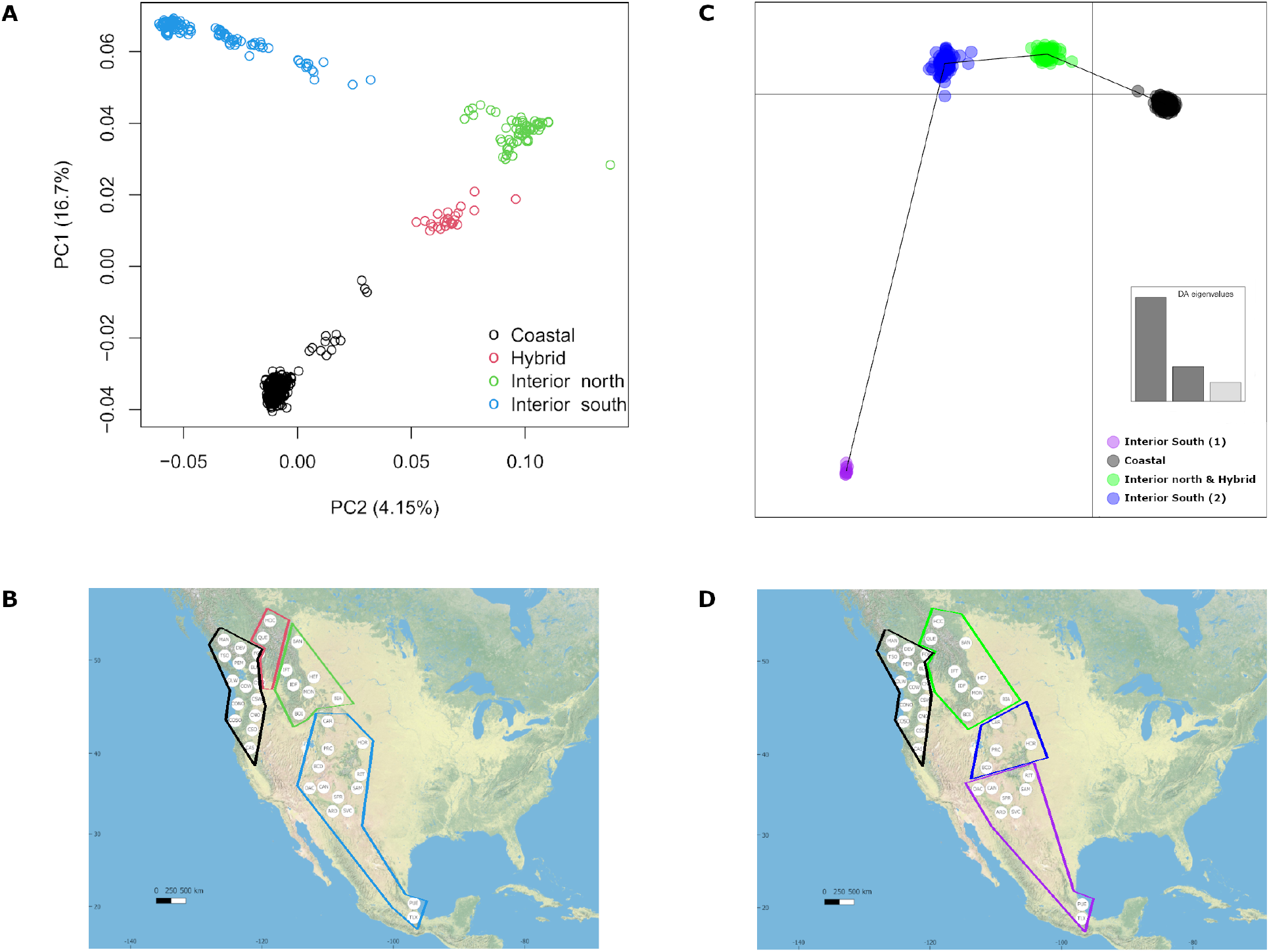
Population genetic structure of Douglas-fir. (A) Principal component analysis of coastal (N=325) and interior (n=188) varieties, including hybrids (n=27), using 11,320 SNPs. (B) Distribution map of populations within PCA clusters. (C) DAPC for Douglas-fir populations. Black lines denote genomic distance between clusters. (D) Distribution map of clusters and populations determined by the DAPC analysis.

Further analysis of the structure of the population with ADMIXTURE resulted in four ancestral populations (best K) and different levels of hybridization across the geographic distribution of the species (Figure 3; Additional file 3; [29]). Each variety was separated into two north to south ancestral populations, with regions of hybridization between them. Coastal Douglas-fir from British Columbia were differentiated from coastal individuals from the United States. Interior individuals from British Columbia (Canada), Idaho, and Montana clustered together, whereas a different genetic cluster was composed by interior individuals in the South (Utah, Colorado, Arizona, New Mexico, Puebla and Tlaxcala-Mexico). The most diverse events of admixture occurred among individuals located in the contact zones in British Columbia and the Washington Cascades (Figure 3). The prevailing admixture component in these individuals was from the interior north variety, suggesting asymmetric introgression from the interior to the coastal variety. The second most important admixture contribution in the hybrids corresponded to the northern ancestral population of the coastal variety in British Columbia, which was coincident with the geographic position of the contact zone. Coastal and interior-north individuals located near the contact zones also presented low levels of admixture among their populations.

**Figure 3.**
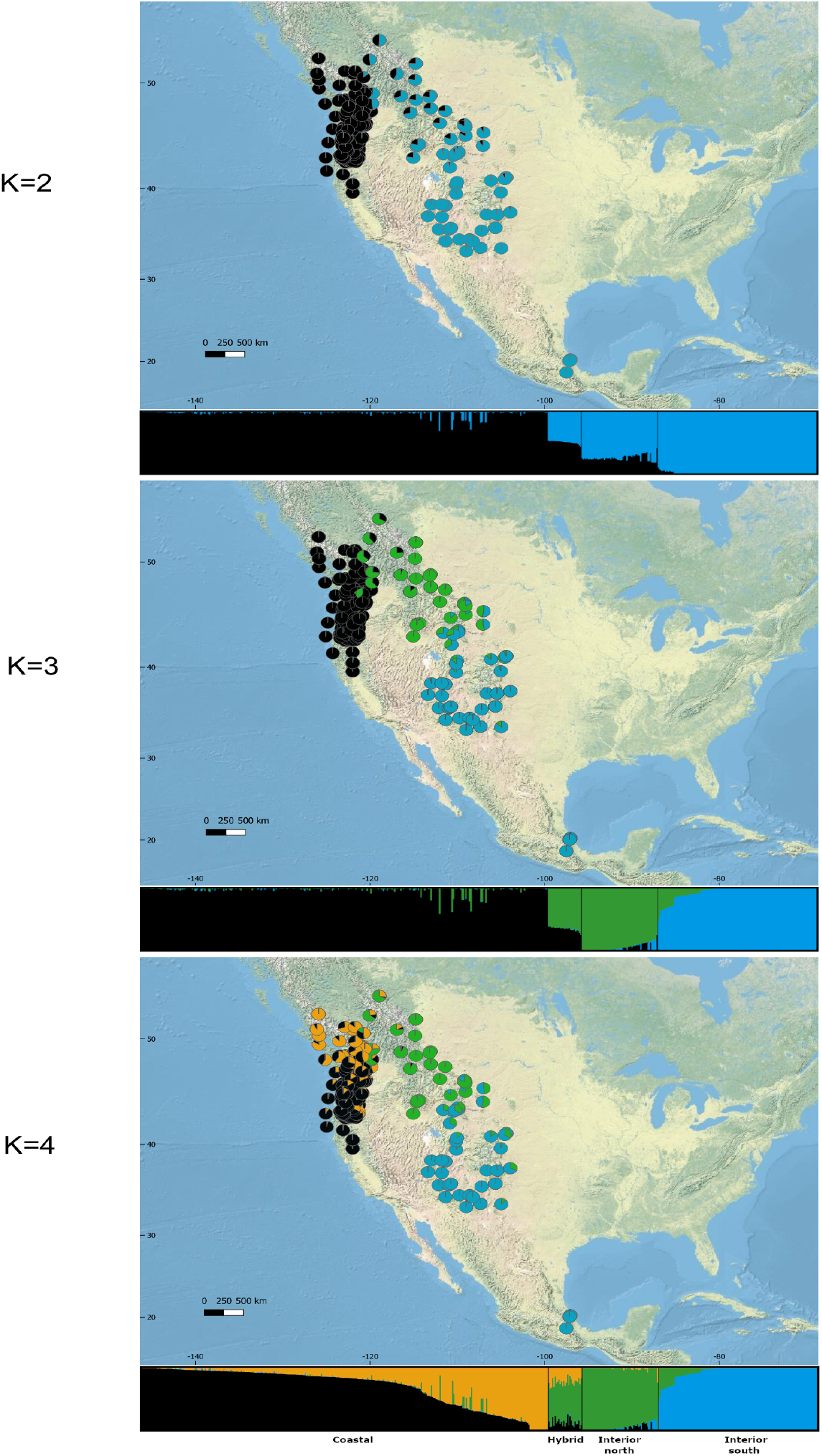
Admixture analysis of Douglas-fir varieties. Clustering of individuals and geographic distribution of admixture proportions obtained with the Admixture software for K2 to K4.

### Hybrid classes

To get further insights into the attributes of the populations and especially the hybrids of the species, the NewHybrids program was used [31]. This program estimates the posterior probability of each individual of the population falling into different hybrid classes. The six genotype groups chosen were: pure interior parent, pure coastal parent, F1 hybrid, F2 hybrid, backcross with pure interior parent, and backcross with pure coastal parent. Almost all coastal and interior-south individuals fell into the pure coastal parent and pure interior parent categories, respectively (Figure 4). The majority (87%) of hybrids were classified as F2s, and no individual was predicted to be a first-generation (F1) hybrid.

**Figure 4.**
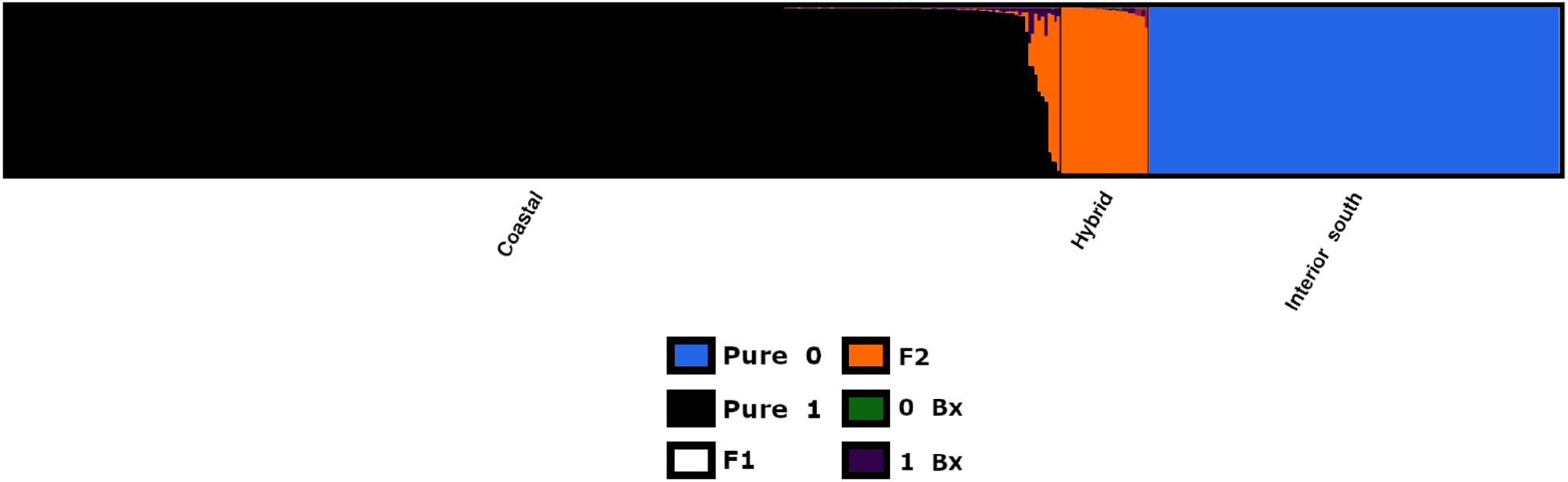
NewHybrids analysis of Douglas-fir trees. Bar plot of posterior probabilities of category membership for each individual of Douglas-fir. Categories are: pure interior parent (Pure 1), pure coastal parent (Pure 0), F1 hybrid (F1), F2 hybrid (F2), backcross with pure interior parent (0 Bx) and backcross with pure coastal parent (1 Bx).

### Genetic diversity, population differentiation, and isolation by distance and environment

Inbreeding, heterozygosity, population differentiation (pairwise Fst) and isolation by distance were estimated from the dataset in this study. The population inbreeding coefficients ranged from 0.08 to 0.92. The highest inbreeding coefficients were found in Mexican populations and southern populations of the interior variety (Figure 5); while the lowest inbreeding coefficients corresponded to coastal and hybrid populations, most of them located in British Columbia (Figure 5). Coincident with inbreeding coefficient values, the lowest heterozygosity values corresponded to the Mexican and interior-south populations (μ=0.09) and the highest values to the coastal and hybrids populations (μ=0.24 and μ=0.23, respectively; Figure 6A). The northern interior populations presented an intermediate mean of 0.17. The two populations with the lowest heterozygosity values corresponded to the populations of the interior variety from Mexico (Puebla and Tlaxcala), while the two populations with the highest values corresponded to coastal populations located in British Columbia (Additional file 4). Higher genetic population differentiation based on pairwise Fst values was observed between populations of the different genetic groups than within their own populations (Figure 6B; Additional file 5). The widest range in Fst values was found when comparing populations of the interior-south genetic group. The coastal and the interior-south populations were among the most genetically differentiated ones. Comparisons involving hybrids showed that these populations were more genetically similar to the interior-north populations, and more genetically distant from the interior-south populations (Figure 6B). Aiming to examine if nearby populations were more genetically similar, a Mantel test was conducted to explore the correlation between geographic and genetic distances. The Mantel test showed a significant strong positive correlation between genetic and geographic distance among all the populations of the species (Figure 6C; r = 0.74 and p = 0.001). Isolation by environment analysis also showed positive correlations between genetic distance and annual mean temperature (Figure 6D; r = 0.35 and p = .0001) and precipitation variables (Figure 6E; r = .11 and p = .0001). A significant positive relationship between geographic distance and temperature (r = .219 and p = .0001) and precipitation (r = .213 and p = .0001) variables was observed as well (Additional file 6).

**Figure 5.**
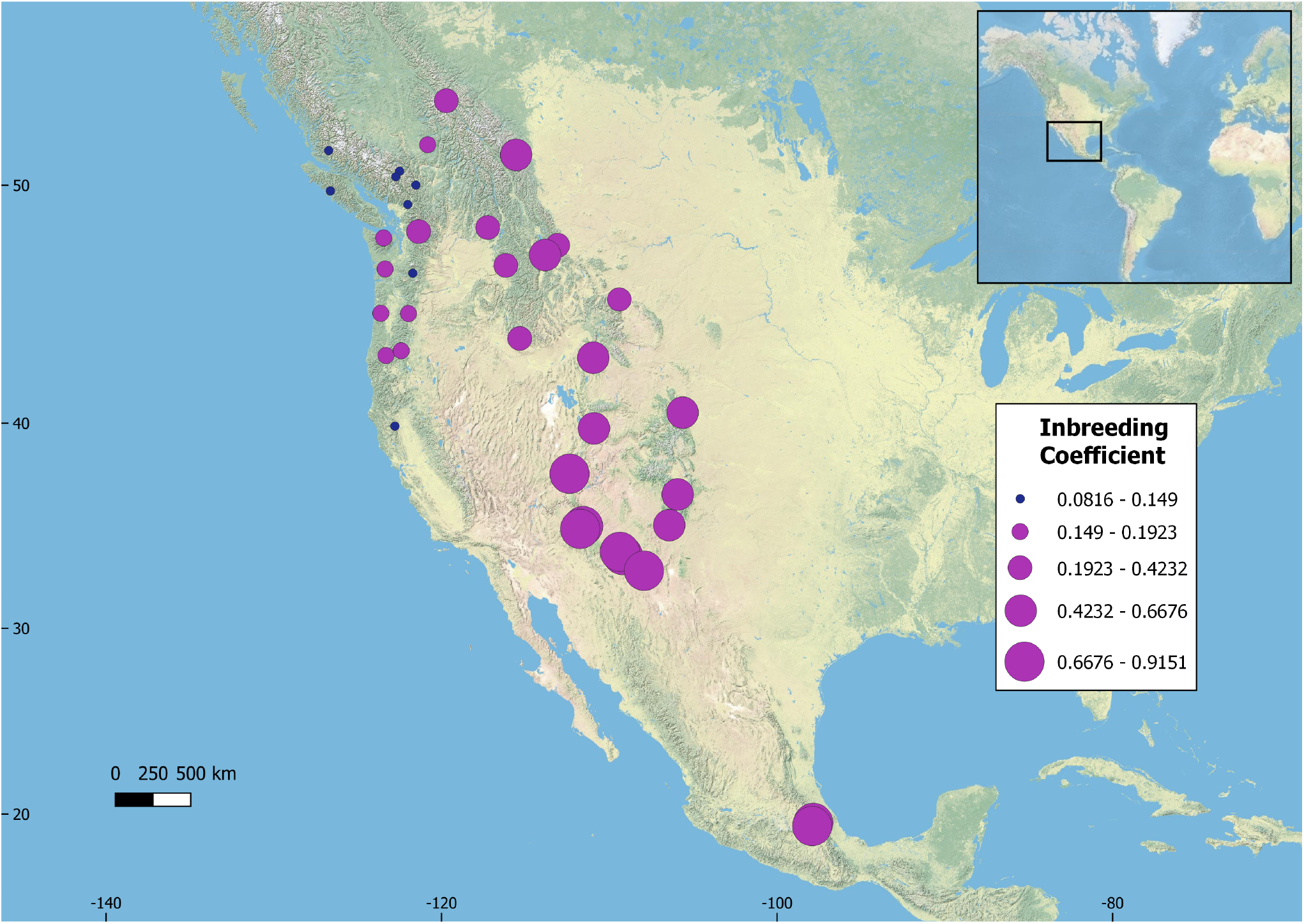
Inbreeding coefficient analysis of Douglas-fir populations. Each circle represents one of the 37 collected populations. Circle size is proportional to the average inbreeding coefficient.

**Figure 6.**
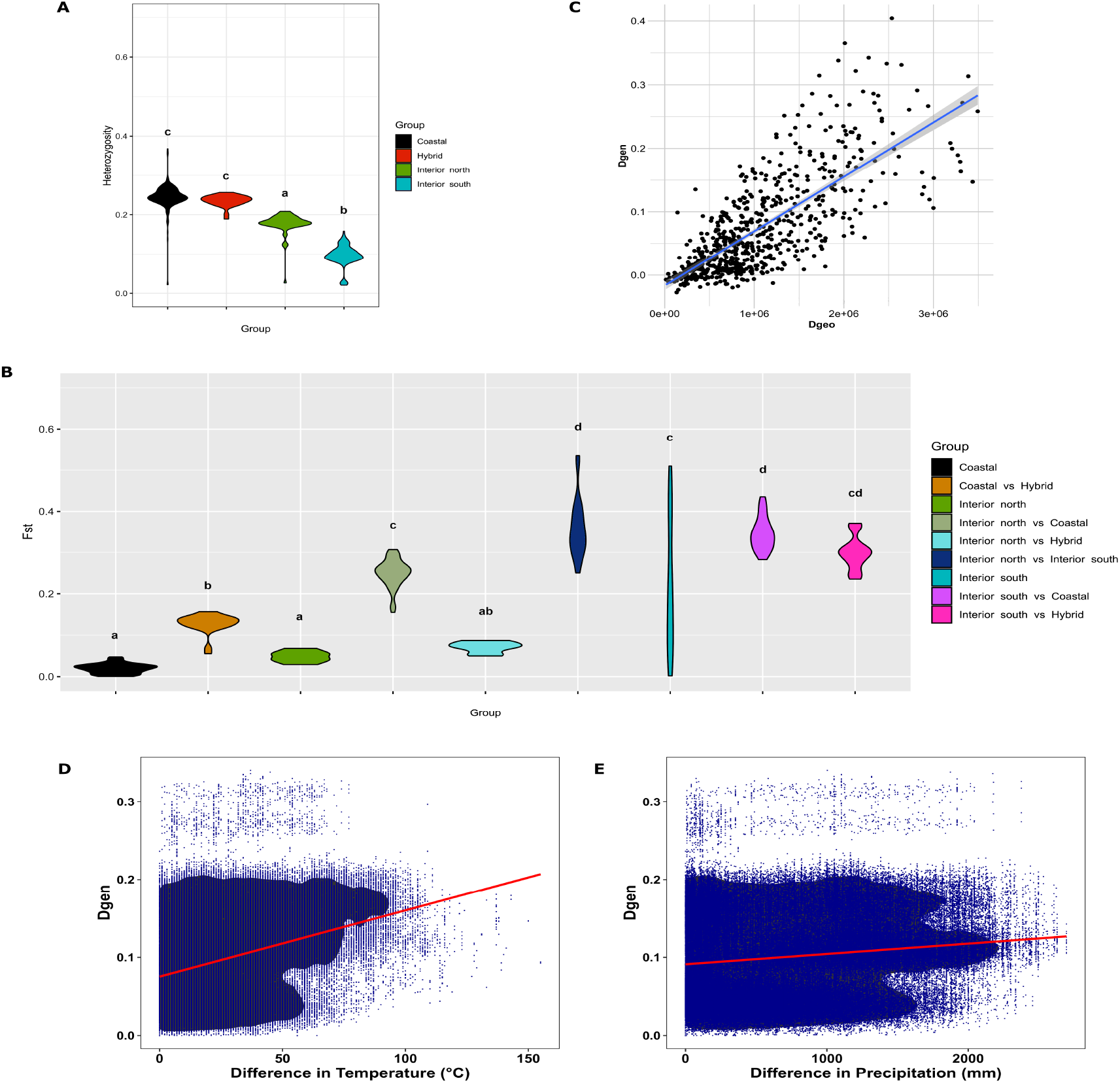
Heterozygosity, Fst and Mantel analyses. Violin plots of heterozygosity (A) and pairwise Fst (B) between coastal, interior-south, interior-north and hybrid groups. (C,D,E) Mantel test analysis of the correlations between geographic, genetic and environmental distances of Douglas-fir populations. Statistical analysis for group comparisons was performed using the SPSS software (version 29.0) through one-way ANOVA followed by Tukey’s test (p ≤ 0.05). Mean values sharing the same letter were not significantly different.

### Signatures of selection

Douglas-fir varieties differ in adaptive traits which allow populations to differentially respond to contrasting environmental conditions. Signatures of selection in loci can reveal evolutionary processes of adaptive differentiation. To detect loci under selection in the Douglas-fir populations, we used BayeScan and pcadapt tools [37,38]. In total, 390 SNPs were identified as outliers by both tools, which matched 300 predicted Douglas-fir genes (Additional file 7). Enrichment analysis based on GO and KEGG pathway terms identified metabolic process, cellular process, and primary metabolism among the most highly represented biological categories. DNA binding, catalytic activity and transcription regulation activity were the top three represented categories in the molecular function classification. For KEEG categories, the most highly represented pathways were phenylalanine metabolism, methane metabolism, and phenylpropanoid biosynthesis.

Signatures of local adaptation can be elucidated by analyzing markers that correlate with climatic data [41]. To evaluate (putative) signatures of local adaptation in Douglas-fir populations, 25 climatic variables related to temperature and precipitation were extracted from ClimateNA database [40]. Bayes factors for all pure and hybrid populations identified 971 unique SNPs associated with the 25 climatic variables. These 971 SNPs were associated with 754 unique predicted genes (Additional file 7). The three most represented KEGG categories for these genes were metabolic metabolism, phenylpropanoid biosynthesis and phenylalanine metabolism. The most represented GO categories for molecular function were catalytic activity, binding, and protein binding. Response to stimulus, response to chemical compounds and metabolic processes were the top three biological processes. The number of SNPs that were associated with environmental variables and under selection were 64 (Figure 7A). These SNPs were assigned to 36 annotated genes (Table 2). In addition, we evaluated associations between genotypes and environments in each variety separately using only the coastal or interior populations (Additional files 8-9). We detected 976 unique SNPs associated with environmental variables in either the interior (596 SNPs) or the coastal populations (380 SNPs). Only 18 SNPs were common between the two varieties (Figure 7B). The top represented GO categories of the genes corresponding to coastal populations for biological processes, cellular component and molecular function were response to stimulus, cell-part, and catalytic activity, respectively. In the case of the interior populations, the top represented GO categories of the genes associated to environmental variables for the biological processes, cellular component and molecular function categories were response to stimulus, cell, and catalytic activity, respectively.

**Figure 7.**
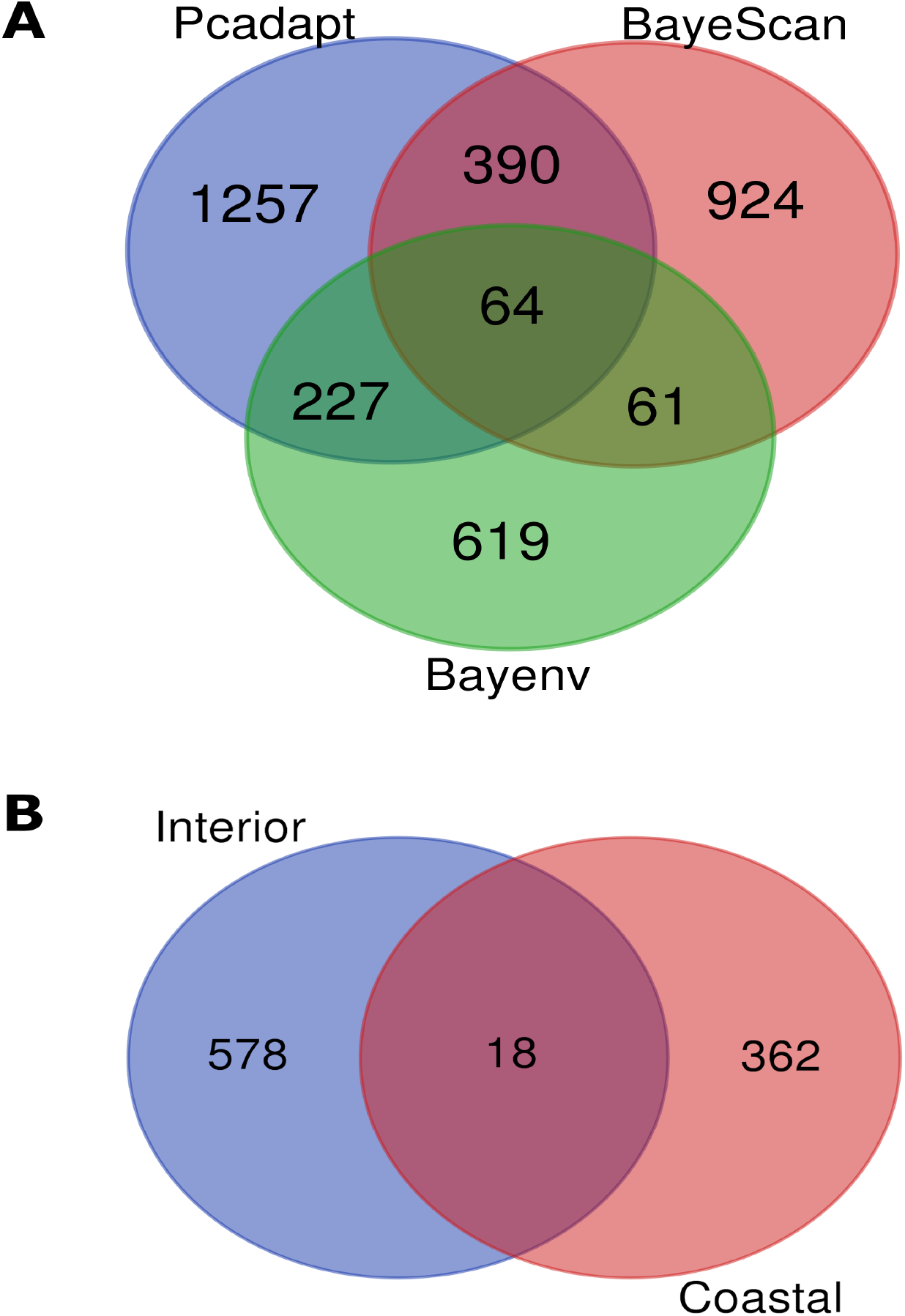
Venn diagrams of the number of outlier SNPs and SNPs-associated with environmental variables. (A) Venn diagram showing overlap between the SNPs detected by each method using all populations of Douglas-fir. (B) Venn diagram showing overlap between the SNPs detected by Bayenv among coastal and interior populations.

**Table 2.**
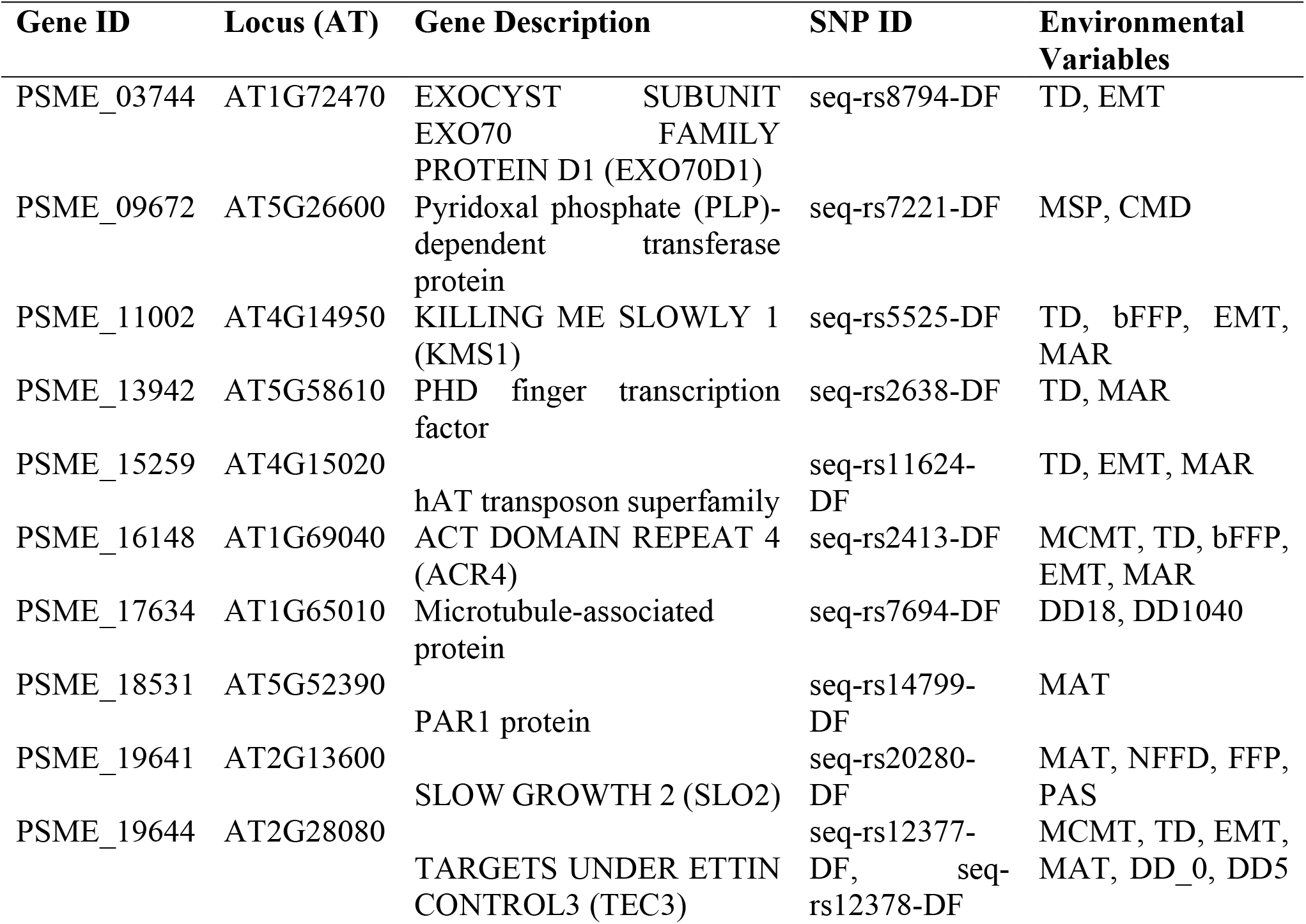
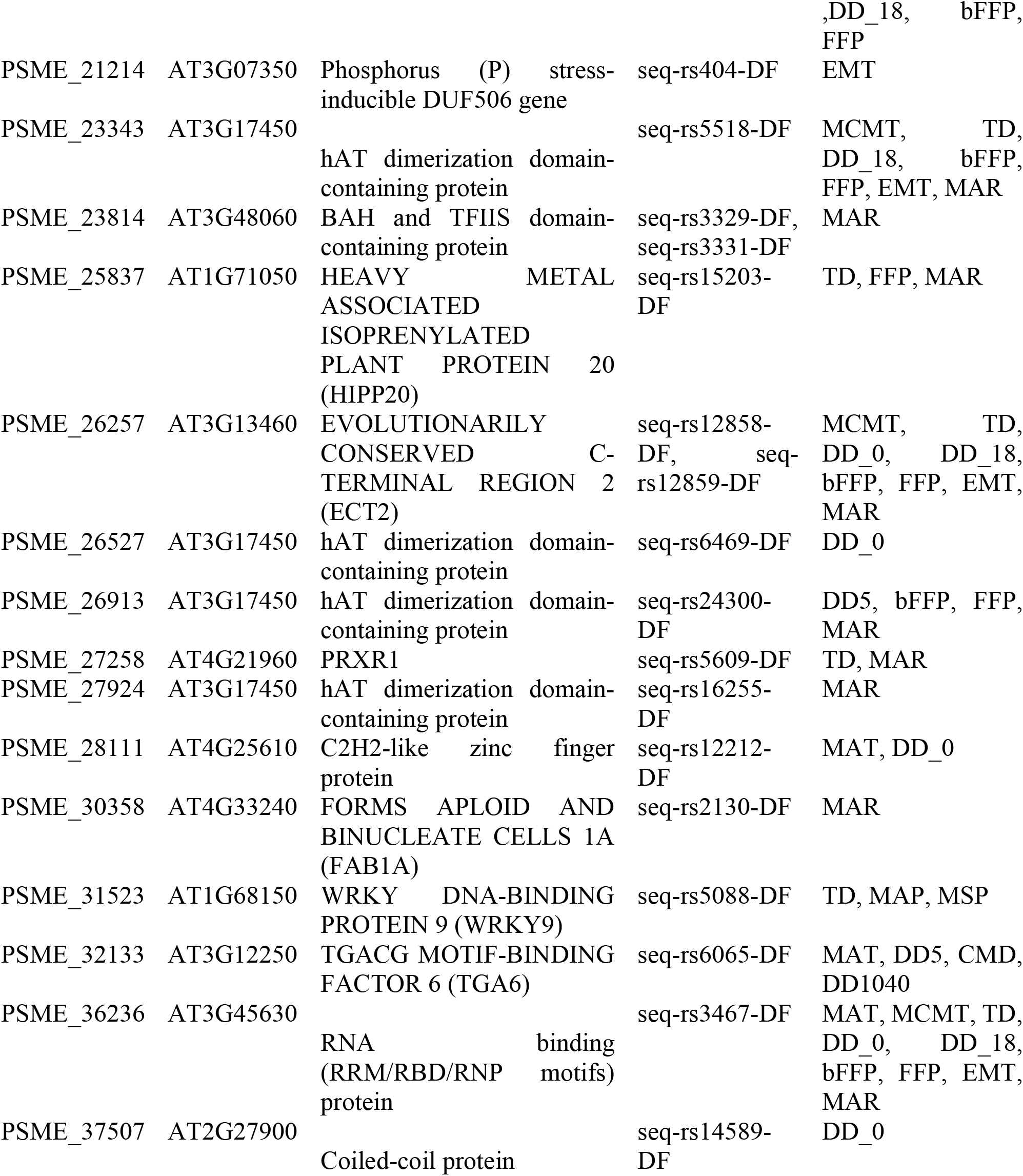
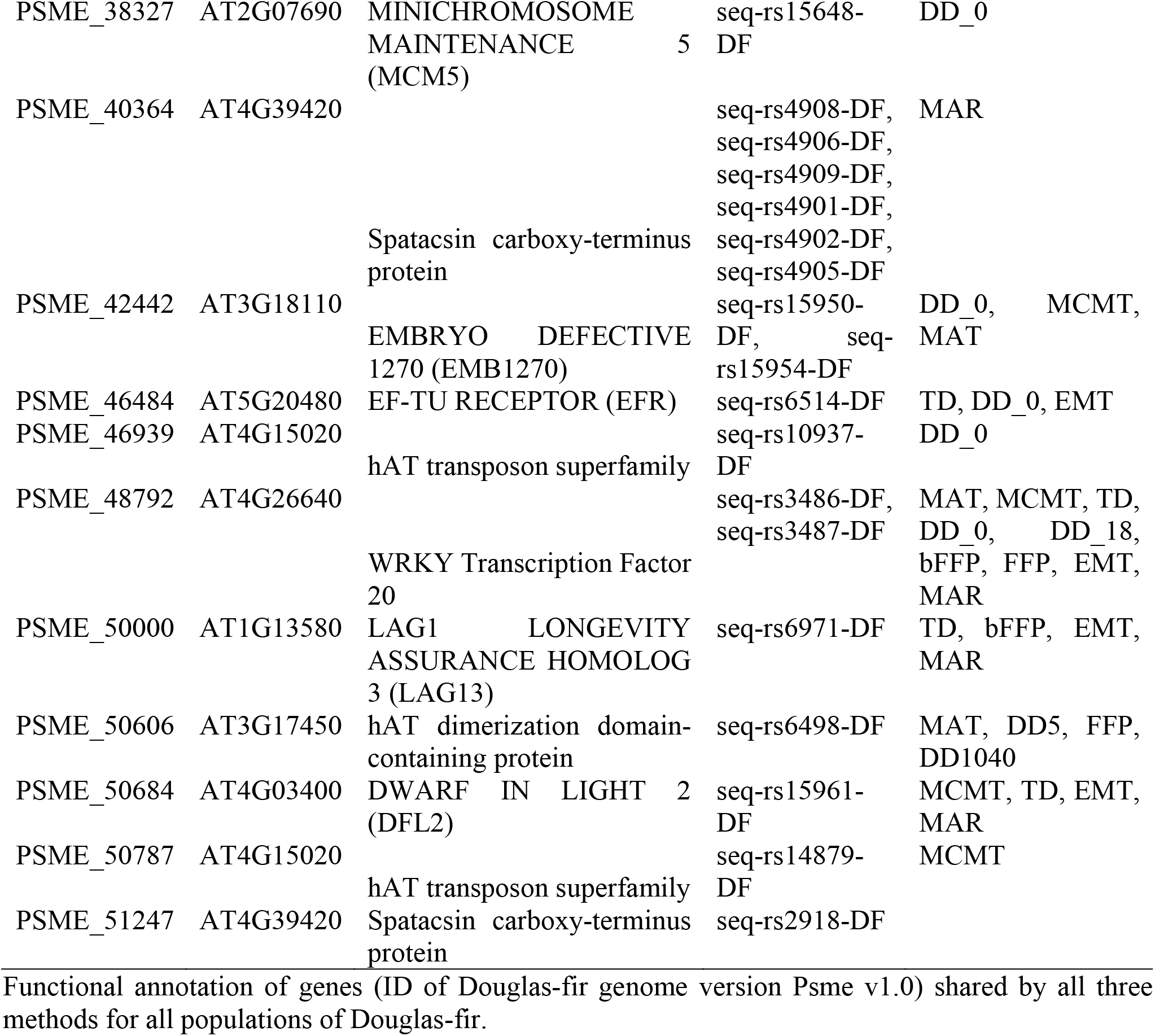
Genes associated with environmental variables and detected as outliers.

### Environmental niche modeling

The characterization of the environmental space occupied by Douglas-fir populations was based on a pairwise comparison that allowed us to identify the climate variables driving a niche differentiation between clusters. Precipitation of the coldest quarter, temperature annual range and maximum temperature of warmest month segregated coastal from interior north populations. Similarly, minimum temperature of the coldest month, mean temperature of the coldest quarter, and precipitation of the coldest quarter separated coastal from hybrid populations. The rest of the relationships are shown in Additional file 10.

The highest bioclimatic niche overlap occurred between interior north–hybrid populations (D=0.26, p=0.05), followed by interior north–interior south (D=0.14, p=0.06) and coastal–interior south (D=0.1, p=0.08). Interestingly, the smallest overlap was found between coastal and both interior north and hybrid populations (D=0.02 and 0.06, respectively), despite coastal being geographically contiguous to both (Additional file 11, Additional file 12A-F). Occurrence density grids, depicting a pairwise comparison of the portion of the environmental space that is more densely occupied by each population, further clarify niche overlap patterns (Additional file 13A-F).

### Projections of potential distribution

Spatially explicit projections of niche models allow us to discern the habitat suitability across the Douglas-Fir range (Figure 8A). For the coastal variety, a large tract of continuous suitable habitat occurs along the Pacific Northwest, especially in Washington, extreme southern British Columbia and Vancouver Island, and Northern California east of the Great Valley. As for interior north populations highest suitability is predicted for most of Idaho (except for the Snake River plain), southeastern British Columbia, central Montana, and eastern portions of Washington and Oregon. Interior south suitability is scattered across the mountain ranges of Colorado, Utah, New Mexico, and Arizona, as well as high elevations in Mexico (Figure 8A).

**Figure 8.**
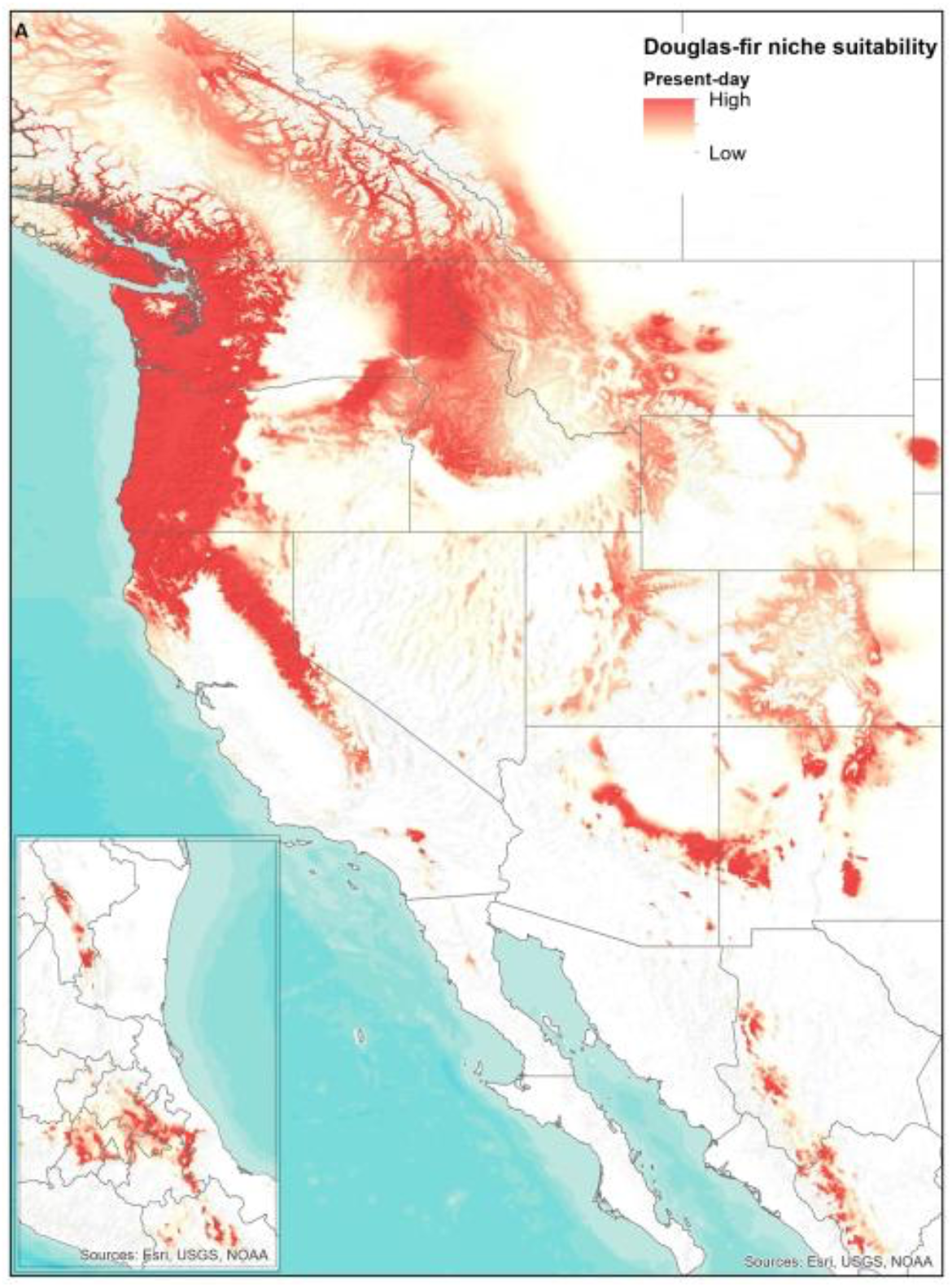
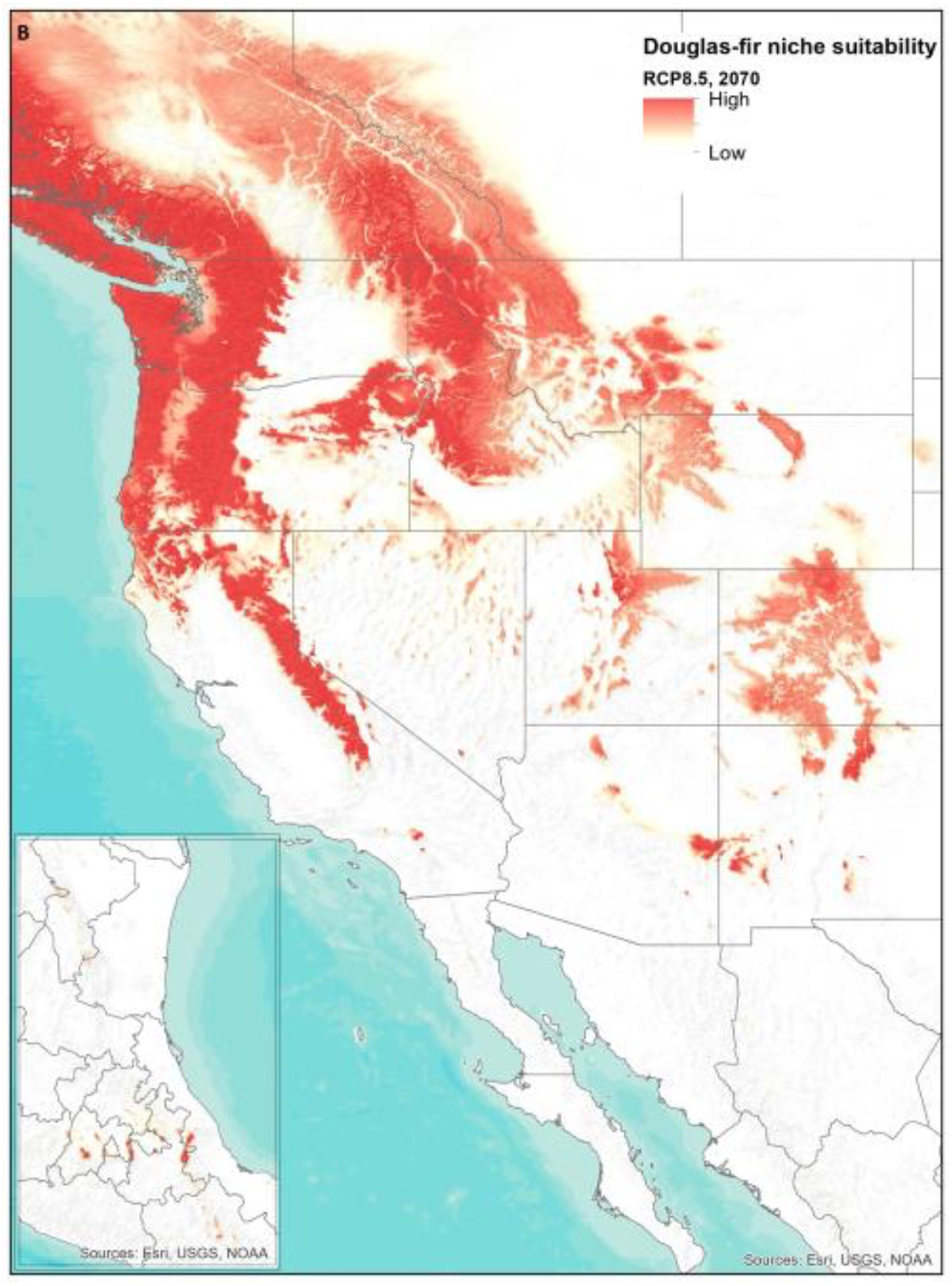
Douglas-fir ecological niche suitability across western North America, with the inset showing eastern Mexico. (A) Present-day potential niche. (B) Potential niche at RCP8.5 scenario at year 2070.

Projections of Douglas-fir potential distribution over the next 50 years showed a leading- and trailing-edge pattern. On one hand, northernmost populations (i.e., coastal, and interior north) are expected to experience a poleward shift with a concomitant overall gain in potential distribution. On the other hand, interior south, and especially Mexican populations are expected to experience severe losses of suitable habitat, particularly at lower elevations, despite some upslope advances on the highest peaks. Depending on any given Representative Concentration Pathway (RCP) scenario between 2050 and 2070, the overall distribution of Douglas-fir in North America could increase by 25-42% (Table 3).

**Table 3.** Douglas-fir potential distribution predicted shifts over the next 50 years in North America.

| Population cluster | Contemporary niche extent (km<sup>2</sup>) | Future niche gains (+) or losses (-) in relation to contemporary niche extent (%) |  |  |  |  |  |  |  |
| --- | --- | --- | --- | --- | --- | --- | --- | --- | --- |
|  |  | RCP2.6 |  | RCP4.5 |  | RCP6.0 |  | RCP8.5 |  |
|  |  | 2050 | 2070 | 2050 | 2070 | 2050 | 2070 | 2050 | 2070 |
| Rangewide | ~ 600,000 | +34 | +33 | +36 | +42 | +34 | +25 | +35 | +32 |
| Coastal | ~ 80,000 | +43 | +30 | +68 | +79 | +79 | +70 | +78 | +110 |
| Interior<br>north | ~ 427,000 | +50 | +46 | +55 | +62 | +62 | +65 | +63 | +78 |
| Interior<br>south | ~ 93,000 | -43 | -41 | -48 | -57 | -42 | -62 | -65 | -78 |

Different Representative Concentration Pathway (RCP) were used.

Nonetheless, on a per-population basis, gains and losses are exacerbated. Under the RCP2.6 scenario in 2070, the coastal population is projected to experience at least a one-third increase in suitable conditions, while in the event of an RCP8.5 scenario during that decade, the increase may exceed a doubling (Table 3). On the contrary, interior south populations are expected to endure a drastic reduction of the contemporary suitable habitat, with losses reaching 65-78% in the worst-case scenarios (Table 3). Changes in suitability for the interior north populations are predicted to be mostly gains although not as pronounced as for those in the Pacific Northwest (Figure 8B). Finally, we consider that small sample size for the hybrid population precluded a robust estimation of their niche shifts across time.

## Discussion

### Role of selection and gene flow maintaining population structure in the species

The environmental adaptation of long-lived trees is crucial for forest survival. The distribution of the species from the genus *Pseudotsuga* is discontinuous, but wide. They are present in North America, Mexico, and Asia. Douglas-fir, the only species from the Pinaceae family with 13 chromosomes, has one of the broadest ranges of any conifer from North America. It has been introduced into temperate regions since the mid-19 century. The natural populations of both varieties of Douglas-fir have high levels of genetic diversity with potential to confer resilience to varying climates. In this study, we used a custom-designed SNP array to study the nuclear genome variation, its relation to space and the local adaptation of several natural populations of Douglas-fir.

The SNPs found in the present study revealed clear genetic structure between the two varieties as previously reported [15,18,19,20,21,22]. The best K-value for the number of ancestral populations in our study was four. The number of ancestral populations previously reported with different genetic markers is variable; however different ancestral populations have been reported within each variety that distinguishes north and south individuals, which indicates that the structure is geographically shaped. Indeed, we observed that Fst values in pairwise comparisons were higher between distant populations of the two varieties, suggesting reduced gene flow in these populations. Strong correlation of patterns of population genetic variation that derive from spatially limited gene flow were also detected through the isolation by distance analysis. Geographic distance was significantly correlated with genetic distance, a stronger correlation was even detected than with environmental variables; however, we also found significant correlation between geographic distance and environmental distances, highlighting spatial and environmental heterogeneous selection in this species.

### Inter-varietal hybridization and introgression

Natural hybridization between the interior and coastal varieties was found in the contact zones in British Columbia and the Washington Cascades. Hybrids in both locations present a higher ancestry from the interior north variety, suggesting asymmetric introgression from the interior to the coastal variety. Asymmetric introgression has been previously found in other conifer species inhabiting the same geographic region in western Canada, such as *Picea glauca* x *P.engelmannii* [49] and *Pinus contorta* x *P. banksiana* [50].

Physical barriers can be important mechanisms in limiting gene flow. Mountain ranges are often a physical barrier to gene flow in widely dispersed species. Conifers, being wind pollinated species, may be susceptible to elevational differences between populations [51]. Douglas-fir grows from sea level to over 3000 m of elevation with northern interior populations growing at the highest altitude [52,8]. Hybrids grow at intermediate elevations between northern coastal and northern interior populations which may explain the higher ancestry of the interior north variety as gene flow from the coastal populations is limited due to the elevational gradient. Previous studies have identified asymmetric introgression in hybrids between *Pinus strobiformis* and *Pinus flexilis* where cold-resilient genes from *P. flexilis* were favored and maintained in hybrid populations [53]. A similar pattern may exist in our study system where asymmetric introgression from interior douglas-fir confers a fitness advantage in natural populations of hybrids. Previous work in douglas-fir suggests that adaptive introgression from the interior variety may have resulted in natural populations of hybrids with increased water-use efficiency (WUE) and heat tolerance [8].

Our results also indicate that the hybrid zones are not of recent origin and that contemporary hybridization between the pure varieties must be rare (as inferred by the lack of F1 hybrids). In contrast, all hybrids found in this study were advanced-generation hybrids, resulting from the cross of two hybrid parents. Backcrosses, in which a hybrid crosses with a pure variety parent were largely absent, suggesting the contact zones are not hybrid swarms. This suggests the hybrid zones are most likely maintained by some level of hybrid fitness, in which hybrids can mate and survive at higher rates than pure varieties in novel environments.

Hybrid individuals may be able to occupy unique novel environments that differ from the environment that the parents experienced. This may occur because of hybridization mechanisms such as transgressive segregation which generates unique packages of alleles and can generate extreme phenotypes that exceed that of the pure varieties [54]. Previous studies have found that transgressive segregation between two ecotypes in *Avena barbata* may contribute to adaptation through the production of novel phenotypes and can result in the colonization of novel niches [55]. Transgressive segregation in hybrid douglas-fir may have allowed establishment of populations in harsh conditions such as the Washington Cascades and mountains in British Columbia. These regions are unique compared to the environments of the pure varieties as they experience high rates of precipitation as snow, low amounts of precipitation, and low mean annual temperature. Genetically distinct hybrid individuals are observed in these harsh environments which may suggest that the advanced generation hybrids that occupy these niches are locally adapted to unique environments compared to the environments of the pure varieties [56]. Future research may include reciprocal transplant studies to examine growth and markers of fitness in common gardens in different environments.

### Candidate loci for local adaptation

Understanding the genomic basis of local adaptation in long-lived trees is crucial to predict their responses to the oncoming effects of climate change. Local adaptation emerges across heterogeneous environments, which as previously shown is the case with Douglas-fir due to its wide distribution. With the use of the Bayenv mixed model we found inter- and intra-varietal differences in the frequency of gene loci that indicate local adaptation in Douglas-fir. The functions of several of the genes associated with environmental variables and detected as outliers are associated with abiotic stress responses and are therefore relevant to local adaptation. For example, we found that some genes (*ECT2*, *SLO2*, and *FAB1A*) that were associated with climatic variables respond or are related to abscisic acid (ABA), which is crucial for abiotic stress responses (Table 2). The ECT2 (EVOLUTIONARILY CONSERVED C-TERMINAL REGION 2) gene encodes a YTH domain-containing reader protein that regulates transcriptional and post-transcriptional gene expression through recognition of m6A modifications. ECT2, ECT3 and ECT4 have genetically redundant functions in ABA response regulation and their disruption destabilizes mRNAs of ABA signaling related genes resulting in ABA hypersensitivity [57]. The SLO2 gene encodes a pentatricopeptide repeat protein that functions as a mitochondrial RNA editing factor. Disruption of SLO2 function results in ABA hypersensitivity, insensitivity to ethylene and increased drought and salt tolerance [58]. The FAB1A gene encodes a 1-phosphatidylinositol-3-phosphate 5-kinase. Null mutants of phosphatidylinositol 3-phosphate 5-kinases in Arabidopsis presented delayed stomatal closure during ABA treatment and increased water loss [59]. Other genes detected as outliers respond to light stimuli (TEC3, BAH and TFIIS domain-containing protein, and DFL2), oxidative stress (PRXR1), and temperature (C2H2-like zinc finger protein). The transcription factor WRKY9 regulates salt tolerance and LAG13 is involved in hypoxia tolerance [60,61].

As probably expected, due to its wide distribution, we also detected signals of local adaptation within populations of each variety in Douglas-fir. Most of the SNPs associated to environmental variables within each variety were not shared, which denotes differentiation and genetic diversity among them; however, the top represented GO term categories in each variety were similar, suggesting that genes associated with environmental variables in both varieties play related roles. Only 18 SNPs were shared between the two varieties. Within the genes that contain shared SNPs, we found genes relevant to stress responses in plants. For example, one of these genes is the previously mentioned SLO2 gene that encodes a pentatricopeptide repeat protein [58]. Also, within this group of genes, we found the gama tonoplast intrinsic protein 2 (*TIP2*) gene, which is involved in ABA and salinity stress responses [62]. The D6 protein kinase like 2 and RAD1, involved in phototropism and resistance to UV radiation, respectively were detected as well [63,64]. An AP2/ERF transcription factor, member of the ethylene signaling pathway, that confers resistance to heat, and hydrogen peroxide stresses was also detected within shared SNPs [65].

### Future distribution of the species under climate change

A few studies have assessed the ecological niche of Douglas-fir, among other conifers, and the potential shifts in distribution in response to future scenarios of climate change [66,67,68]. In general, the evidence we found in our study supports the general findings of those investigations, which can be summarized in two major aspects. First, the ample separation regarding the environmental space occupied by the two main Douglas-fir varieties (i.e., coastal, inland), which in turn is reflected in a geographic clustering on a regional scale, coherent with their genetic structuring; and second, the contrasting impacts of climate change over the next decades on the distribution of the distinct populations. As global climate gets increasingly warmer, leading-edge populations of temperate tree species such as Douglas-fir are expected to migrate poleward [69,70,71], either to track suitable conditions or to colonize new available areas, while trailing-edge populations will be forced to engage in upslope migrations, which nonetheless will be greatly constrained by local geomorphological features. As a result, over the next 50 years, the Douglas-fir suitable habitat is expected to increase at least 25% in relation to their current extent. The potential gains are even larger when considering the extreme warming that could take place in an RCP8.5 scenario. Nevertheless, the considered timespan (5 decades) is certainly small considering the longevity and generation time of the Douglas-fir. Beyond that period, erratic oscillations in global climate patterns could disrupt any temporary equilibrium in ecological communities, including temperate coniferous forests.

## Conclusions

Our range-wide population structure and genetic diversity analyses of Douglas-fir varieties indicate high genetic variation in their populations with clear structured populations and differential adaptive potential to different climate scenarios as result of spatially heterogenous selection and dissimilar evolutionary histories. The genetic variation associated with climate data found here helps our understanding of the molecular mechanisms controlling long-living trees adaptation. The future distributions predicted in this study revealed different evolutionary paths for Douglas-fir varieties, which constitutes valuable information regarding the conservation and management of the species.

## Declarations

### Ethics approval and consent to participate

Not applicable

### Consent for publication

Not applicable

### Availability of data and materials

The attached supplementary files contains the data.

## Funding

This project was supported by the National Science Foundation [CAREER project 2145834] to A.R.D.L.T.

## Competing interests

The authors declare no competing interests.

## Authors’ Contributions

A.R.D.L.T. designed and supervised the project and obtained research funding; K.B. and J.R.M collected samples; K.B performed all lab analyses; P.P. carried out all population genomic analyses; G.P.L. performed the climate niche modeling; P.P., A.R.D.L.T, K.B. and G.P.L. wrote the manuscript; all authors approved the final version of the manuscript.

## Acknowledgements

The authors would like to thank Benjamin Wilhite and Matthew Weiss for collecting some of the samples used in this study.

## Additional files

**Additional file 1. Environmental variables for SNP-climate associations.**

File name: Additional file 4.xlsx

**Additional file 2. Summary of sample collection.**

File name: Additional file 4.xlsx

**to K4.**

**Additional file 3. Bar plots of admixture proportions of Douglas-fir populations from K2**

File name: Additional file 3.TIFF

**Additional file 4. Violin plots of heterozygosity values per population.**

File name: Additional file 4.TIFF

**Additional file 5. Pairwise Fst values per population.**

File name: Additional file 5.xlsx

**Additional file 6. Mantel test correlations between geographic and environmental distances.**

File name: Additional file 6.TIFF

**Additional file 7. SNPs associated with environmental variables and detected as outliers in Douglas-fir populations.**

File name: Additional file 7.xlsx

**Additional file 8. SNPs associated with environmental variables of coastal populations.**

File name: Additional file 8.xlsx

**Additional file 9. SNPs associated with environmental variables of interior populations.**

File name: Additional file 9.xlsx

**Additional file 10. Environmental ordination of Douglas-fir populations based on Principal Component Analysis characterization of their Hutchinsonian niche.**

File name: Additional file 10.docx

**Additional file 11. Pairwise niche overlap (D) between Douglas-fir population clusters. P-values are shown below the diagonal.**

File name: Additional file 11.docx

**Additional file 12. Niche overlap (D) between Douglas-fir populations.** (A) Coastal-Hybrid.

(B) Coastal-Interior north. (C) Coastal-Interior south. (D) Hybrid-Interior north. (E) Hybrid-Interior south. (F). Interior north-Interior south.

File name: Additional file 12.docx

**Additional file 13. Pairwise comparison of occurrence density grids.** (A) Coastal-Hybrid.

(B) Coastal-Interior north. (C) Coastal-Interior south. (D) Hybrid-Interior north. (E) Hybrid-Interior south. (F). Interior north-Interior south. File name: Additional file 13.docx

## Notes

### Competing Interest Statement

The authors have declared no competing interest.

## References

1. Sork VL, Aitken SN, Dyer RJ, Eckert AJ, Legendre P, Neale DB. Putting the landscape into the genomics of trees: approaches for understanding local adaptation and population responses to changing climate. Tree Genet Genomes. 2013;9:901–11.

2. Savolainen O, Pyhäjärvi T, Knürr T. Gene Flow and Local Adaptation in Trees. Annu Rev Ecol Evol Syst. 2007;38:595–619.

3. Feng L, Du FK. Landscape Genomics in Tree Conservation Under a Changing Environment. Front Plant Sci. 2022;13.

4. Bansal S, Harrington CA, Gould PJ, St.Clair JB. Climate-related genetic variation in drought-resistance of Douglas-fir (Pseudotsuga menziesii). Glob Chang Biol. 2015;21:947–58.

5. Montwé D, Spiecker H, Hamann A. Five decades of growth in a genetic field trial of Douglas-fir reveal trade-offs between productivity and drought tolerance. Tree Genet Genomes. 2015;11:29.

6. Marias DE, Meinzer FC, Woodruff DR, McCulloh KA. Thermotolerance and heat stress responses of Douglas-fir and ponderosa pine seedling populations from contrasting climates. Tree Physiol. 2016. 10.1093/treephys/tpw117.

7. De La Torre AR, Wilhite B, Puiu D, St. Clair JB, Crepeau MW, Salzberg SL, et al. Dissecting the Polygenic Basis of Cold Adaptation Using Genome-Wide Association of Traits and Environmental Data in Douglas-fir. Genes (Basel). 2021;12:110.

8. Compton S, Stackpole C, Dixit A, Sekhwal MK, Kolb T, De la Torre AR. Differences in heat tolerance, water use efficiency and growth among Douglas-fir families and varieties evidenced by GWAS and common garden studies. AoB Plants. 2023;15.

9. Domínguez-Álvarez FA. Análisis histórico-ecológico de los bosques de Pseudotsuga en México. Folleto Técnico INIFAP. 1994:43.

10. Mosseler A, Major JE, Simpson JD, Daigle B, Lange K, Park Y-S, et al. Indicators of population viability in red spruce, Picea rubens. I. Reproductive traits and fecundity. Canadian Journal of Botany. 2000;78:928–40.

11. Mápula-larreta M, López-Upton J, Vargas-Hernández JJ, Hernández-Livera A. Reproductive indicators in natural populations of Douglas-fir in Mexico. Biodivers Conserv. 2007;16:727–42.

12. Reyes-Hernández V, Vargas-Hernández J, López-Upton J, Vaquera-Huerta H. Phenotypic similarity among Mexican populations of Pseudotsuga Carr. Agrociencia. 2006;40:545–56.

13. Acevedo-Rodriguez R, Vargas-Hernandez JJ, Lopez-Upton J, Mendoza JV. Effect of geographic origin and nutrition on shoot prenology of Mexican Douglas-Fir (Pseudotsuga sp.) seedlings. Agrociencia. 2006;40:125–37.

14. Cruz-Nicolás J, Vargas-Hernández JJ, Ramírez-Vallejo P, López-Upton J. Genetic diversity and differentiation of Pseudotsuga menziesii (Mirb.) Franco populations in Mexico. Revista fitotecnia Mexicana. 2011;34:233–40.

15. Gugger PF, Sugita S, Cavender-Bares J. Phylogeography of Douglas-fir based on mitochondrial and chloroplast DNA sequences: testing hypotheses from the fossil record. Mol Ecol. 2010;19:1877– 97.

16. López-Upton J, Valdez-Lazalde J, Ventura-Ríos A, Vargas-Hernández J, Guerra-de-la-Cruz V. Extinction Risk of Pseudotsuga Menziesii Populations in the Central Region of Mexico: An AHP Analysis. Forests. 2015;6:1598–612.

17. Wehenkel C, Mariscal-Lucero S del R, Jaramillo-Correa JP, López-Sánchez CA, Vargas-Hernández JJ, Sáenz-Romero C. Genetic Diversity and Conservation of Mexican Forest Trees. 2017. p. 37–67.

18. van Loo M, Hintsteiner W, Pötzelsberger E, Schüler S, Hasenauer H. Intervarietal and intravarietal genetic structure in Douglas-fir: nuclear SSRs bring novel insights into past population demographic processes, phylogeography, and intervarietal hybridization. Ecol Evol. 2015;5:1802–17.

19. Wei X-X, Beaulieu J, Khasa DP, Vargas-Hernández J, López-Upton J, Jaquish B, et al. Range-wide chloroplast and mitochondrial DNA imprints reveal multiple lineages and complex biogeographic history for Douglas-fir. Tree Genet Genomes. 2011;7:1025–40.

20. Li P, Adams WT. Range-wide patterns of allozyme variation in Douglas-fir (Pseudotsuga menziesii). Canadian Journal of Forest Research. 1989;19:149–61.

21. Neophytou C, Weisser A-M, Landwehr D, Šeho M, Kohnle U, Ensminger I, et al. Assessing the relationship between height growth and molecular genetic variation in Douglas-fir (Pseudotsuga menziesii) provenances. Eur J For Res. 2016;135:465–81.

22. Hintsteiner WJ, van Loo M, Neophytou C, Schueler S, Hasenauer H. The geographic origin of old Douglas-fir stands growing in Central Europe. Eur J For Res. 2018;137:447–61.

23. Eckert AJ, Wegrzyn JL, Pande B, Jermstad KD, Lee JM, Liechty JD, et al. Multilocus Patterns of Nucleotide Diversity and Divergence Reveal Positive Selection at Candidate Genes Related to Cold Hardiness in Coastal Douglas Fir (Pseudotsuga menziesii var. menziesii). Genetics. 2009;183:289–98.

24. Müller T, Freund F, Wildhagen H, Schmid KJ. Targeted re-sequencing of five Douglas-fir provenances reveals population structure and putative target genes of positive selection. Tree Genet Genomes. 2015;11:816.

25. Hess M, Wildhagen H, Junker LV, Ensminger I. Transcriptome responses to temperature, water availability and photoperiod are conserved among mature trees of two divergent Douglas-fir provenances from a coastal and an interior habitat. BMC Genomics. 2016;17:682.

26. Nelson TC, Stathos AM, Vanderpool DD, Finseth FR, Yuan Y, Fishman L. Ancient and recent introgression shape the evolutionary history of pollinator adaptation and speciation in a model monkeyflower radiation (Mimulus section Erythranthe). PLoS Genet. 2021;17:e1009095.

27. Zheng X, Levine D, Shen J, Gogarten SM, Laurie C, Weir BS. A high-performance computing toolset for relatedness and principal component analysis of SNP data. Bioinformatics. 2012;28:3326– 8.

28. Jombart T. adegenet: a R package for the multivariate analysis of genetic markers. Bioinformatics. 2008;24:1403–5.

29. Alexander DH, Novembre J, Lange K. Fast model-based estimation of ancestry in unrelated individuals. Genome Res. 2009;19:1655–64.

30. Kopelman NM, Mayzel J, Jakobsson M, Rosenberg NA, Mayrose I. Clumpak: a program for identifying clustering modes and packaging population structure inferences across K. Mol Ecol Resour. 2015;15:1179–91.

31. Anderson EC, Thompson EA. A Model-Based Method for Identifying Species Hybrids Using Multilocus Genetic Data. Genetics. 2002;160:1217–29.

32. Ochoa A, Storey JD. Estimating FST and kinship for arbitrary population structures. PLoS Genet. 2021;17:e1009241.

33. Little EL. Atlas of United States trees. Washington, D.C: U.S. Dept. of Agriculture, Forest Service; 1971.

34. Danecek P, Auton A, Abecasis G, Albers CA, Banks E, DePristo MA, et al. The variant call format and VCFtools. Bioinformatics. 2011;27:2156–8.

35. Wickham H. ggplot2. New York, NY: Springer New York; 2009.

36. Gruber B, Unmack PJ, Berry OF, Georges A. dartr: An r package to facilitate analysis of SNP data generated from reduced representation genome sequencing. Mol Ecol Resour. 2018;18:691–9.

37. Foll M, Gaggiotti O. A Genome-Scan Method to Identify Selected Loci Appropriate for Both Dominant and Codominant Markers: A Bayesian Perspective. Genetics. 2008;180:977–93.

38. Luu K, Bazin E, Blum MGB. pcadapt: an R package to perform genome scans for selection based on principal component analysis. Mol Ecol Resour. 2017;17:67–77.

39. Bu D, Luo H, Huo P, Wang Z, Zhang S, He Z, et al. KOBAS-i: intelligent prioritization and exploratory visualization of biological functions for gene enrichment analysis. Nucleic Acids Res. 2021;49:W317–25.

40. Wang T, Hamann A, Spittlehouse D, Carroll C. Locally Downscaled and Spatially Customizable Climate Data for Historical and Future Periods for North America. PLoS One. 2016;11:e0156720.

41. Günther T, Coop G. Robust Identification of Local Adaptation from Allele Frequencies. Genetics. 2013;195:205–20.

42. Kass JM, Pinilla-Buitrago GE, Paz A, Johnson BA, Grisales Betancur V, Meenan SI, et al. wallace 2: a shiny app for modeling species niches and distributions redesigned to facilitate expansion via module contributions. Ecography. 2023;2023.

43. Phillips SJ, Anderson RP, Schapire RE. Maximum entropy modeling of species geographic distributions. Ecol Modell. 2006;190:231–59.

44. Phillips SJ, Anderson RP, Dudík M, Schapire RE, Blair ME. Opening the black box: an open source release of Maxent. Ecography. 2017;40:887–93.

45. Hijmans RJ, Cameron SE, Parra JL, Jones PG, Jarvis A. Very high resolution interpolated climate surfaces for global land areas. International Journal of Climatology. 2005;25:1965–78.

46. Hutchinson GE. Concluding Remarks. Cold Spring Harb Symp Quant Biol. 1957;22:415–27.

47. Schoener TW. The Anolis Lizards of Bimini: Resource Partitioning in a Complex Fauna. Ecology. 1968;49:704–26.

48. Elith J, Phillips SJ, Hastie T, Dudík M, Chee YE, Yates CJ. A statistical explanation of MaxEnt for ecologists. Divers Distrib. 2011;17:43–57.

49. De La Torre AR, Roberts DR, Aitken SN. Genome-wide admixture and ecological niche modelling reveal the maintenance of species boundaries despite long history of interspecific gene flow. Mol Ecol. 2014;23:2046–59.

50. Yaremchuk D. Differential Introgression Between the Northern and Southern Extents of the Lodgepole x Jack Pine Hybrid Zone Suggests Environmentally-driven Selection and Local Adaptation. Carleton University; 2023.

51. Li Y-S, Shih K-M, Chang C-T, Chung J-D, Hwang S-Y. Testing the Effect of Mountain Ranges as a Physical Barrier to Current Gene Flow and Environmentally Dependent Adaptive Divergence in Cunninghamia konishii (Cupressaceae). Front Genet. 2019;10.

52. Gould PJ, Harrington CA, St. Clair JB. Incorporating genetic variation into a model of budburst phenology of coast Douglas-fir (Pseudotsuga menziesii var. menziesii). Canadian Journal of Forest Research. 2011;41:139–50.

53. Menon M, Bagley JC, Page GFM, Whipple A V., Schoettle AW, Still CJ, et al. Adaptive evolution in a conifer hybrid zone is driven by a mosaic of recently introgressed and background genetic variants. Commun Biol. 2021;4:160.

54. Rieseberg LH, Widmer A, Arntz AM, Burke B. The genetic architecture necessary for transgressive segregation is common in both natural and domesticated populations. Philos Trans R Soc Lond B Biol Sci. 2003;358:1141–7.

55. Johansen-Morris AD, Latta RG. FITNESS CONSEQUENCES OF HYBRIDIZATION BETWEEN ECOTYPES OF AVENA BARBATA: HYBRID BREAKDOWN, HYBRID VIGOR, AND TRANSGRESSIVE SEGREGATION. Evolution (N Y). 2006;60:1585–95.

56. Kagawa K, Takimoto G. Hybridization can promote adaptive radiation by means of transgressive segregation. Ecol Lett. 2018;21:264–74.

57. Song P, Wei L, Chen Z, Cai Z, Lu Q, Wang C, et al. m6A readers ECT2/ECT3/ECT4 enhance mRNA stability through direct recruitment of the poly(A) binding proteins in Arabidopsis. Genome Biol. 2023;24:103.

58. Zhu Q, Dugardeyn J, Zhang C, Mühlenbock P, Eastmond PJ, Valcke R, et al. The Arabidopsis thaliana RNA Editing Factor SLO2, which Affects the Mitochondrial Electron Transport Chain, Participates in Multiple Stress and Hormone Responses. Mol Plant. 2014;7:290–310.

59. Bak G, Lee E-J, Lee Y, Kato M, Segami S, Sze H, et al. Rapid Structural Changes and Acidification of Guard Cell Vacuoles during Stomatal Closure Require Phosphatidylinositol 3,5-Bisphosphate. Plant Cell. 2013;25:2202–16.

60. Krishnamurthy P, Vishal B, Bhal A, Kumar PP. WRKY9 transcription factor regulates cytochrome P450 genes CYP94B3 and CYP86B1, leading to increased root suberin and salt tolerance in Arabidopsis. Physiol Plant. 2021;172:1673–87.

61. Xie L-J, Chen Q-F, Chen M-X, Yu L-J, Huang L, Chen L, et al. Unsaturation of Very-Long-Chain Ceramides Protects Plant from Hypoxia-Induced Damages by Modulating Ethylene Signaling in Arabidopsis. PLoS Genet. 2015;11:e1005143.

62. Pih KT, Kabilan V, Lim JH, Kang SG, Piao HL, Jin JB, et al. Characterization of two new channel protein genes in Arabidopsis. Mol Cells. 1999;9:84–90.

63. Willige BC, Ahlers S, Zourelidou M, Barbosa ICR, Demarsy E, Trevisan M, et al. D6PK AGCVIII Kinases Are Required for Auxin Transport and Phototropic Hypocotyl Bending in Arabidopsis. Plant Cell. 2013;25:1674–88.

64. Jiang C-Z, Yen C-N, Cronin K, Mitchell D, Britt AB. UV- and Gamma-Radiation Sensitive Mutants of Arabidopsis thaliana. Genetics. 1997;147:1401–9.

65. Ogawa T, Pan L, Kawai-Yamada M, Yu L-H, Yamamura S, Koyama T, et al. Functional Analysis of Arabidopsis Ethylene-Responsive Element Binding Protein Conferring Resistance to Bax and Abiotic Stress-Induced Plant Cell Death. Plant Physiol. 2005;138:1436–45.

66. Rehfeldt GE, Jaquish BC, López-Upton J, Sáenz-Romero C, St Clair JB, Leites LP, et al. Comparative genetic responses to climate for the varieties of Pinus ponderosa and Pseudotsuga menziesii: Realized climate niches. For Ecol Manage. 2014;324:126–37.

67. Campbell JL, Shinneman DJ. Potential influence of wildfire in modulating climate-induced forest redistribution in a central Rocky Mountain landscape. Ecol Process. 2017;6:7.

68. Zhao Y, O’Neill GA, Wang T. Predicting fundamental climate niches of forest trees based on species occurrence data. Ecol Indic. 2023;148:110072.

69. Corlett RT, Westcott DA. Will plant movements keep up with climate change? Trends Ecol Evol. 2013;28:482–8.

70. Périé C, de Blois S. Dominant forest tree species are potentially vulnerable to climate change over large portions of their range even at high latitudes. PeerJ. 2016;4:e2218.

71. Elsen PR, Saxon EC, Simmons BA, Ward M, Williams BA, Grantham HS, et al. Accelerated shifts in terrestrial life zones under rapid climate change. Glob Chang Biol. 2022;28:918–35.

