## Supplemental file 10 for "Spatially heterogeneous selection and inter-varietal differentiation maintain population structure and local adaptation in a widespread conifer"

| Additional file 10. Environmental ordination of Douglas-fir populations based on a Principal Component Analysis characterization of their Hutchinsonian niche | | | |
| --- | --- | --- | --- |
| Groups | Variables with highest absolute loadings | PC1 (%) | PC2 (%) |
| Coastal – Hybrid | BIO6, BIO11, BIO17 | 42.5 | 38.1 |
| Coastal – Interior north | BIO19, BIO7, BIO5 | 43.9 | 33.5 |
| Coastal – Interior south | BIO1, BIO11, BIO13 | 39.8 | 35.3 |
| Hybrid – Interior north | BIO7, BIO19, BIO1 | 41.6 | 32.8 |
| Hybrid – Interior south | BIO11, BIO1, BIO12 | 46.6 | 29.3 |
| Interior north – Interior south | BIO11, BIO6, BIO12 | 43.7 | 29.3 |
