## Supplemental file 11 for "Spatially heterogeneous selection and inter-varietal differentiation maintain population structure and local adaptation in a widespread conifer"

| Additional file 11. Pairwise niche overlap (D) between Douglas-fir population clusters. *P*-values are shown below the diagonal | | | | |
| --- | --- | --- | --- | --- |
|  | Coastal | Hybrid | Interior north | Interior south |
| Coastal | -- | 0.06 | 0.02 | 0.1 |
| Hybrid | 0.19 | -- | 0.26 | 0.09 |
| Interior north | 0.37 | 0.05 | -- | 0.14 |
| Interior south | 0.08 | 0.02 | 0.06 | -- |
