## Supplementary figures and images for "Spatially heterogeneous selection and inter-varietal differentiation maintain population structure and local adaptation in a widespread conifer"

### Supplemental file 3

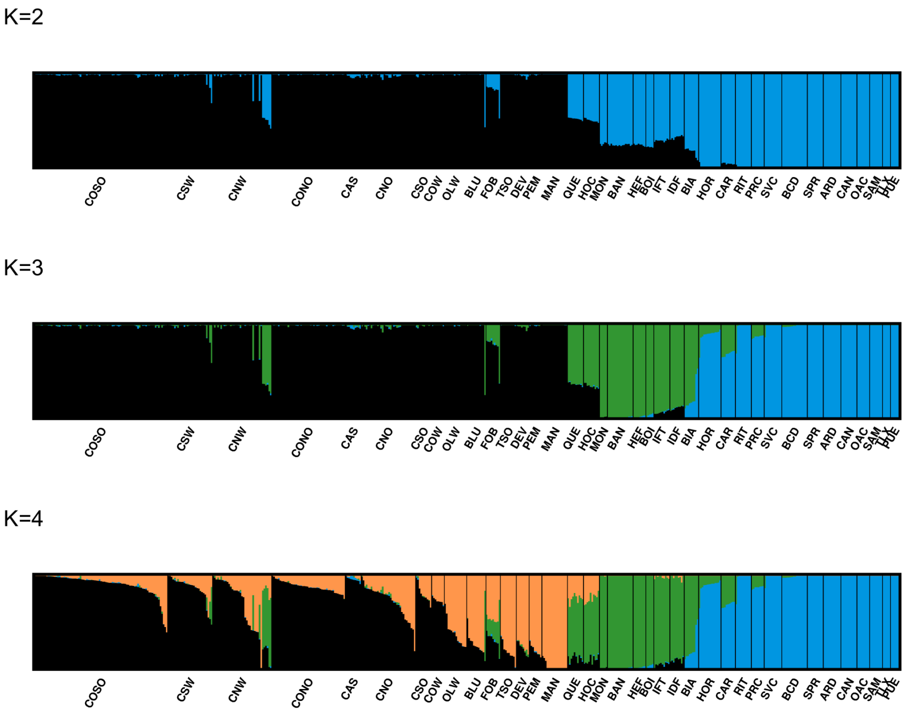

### Supplemental file 4

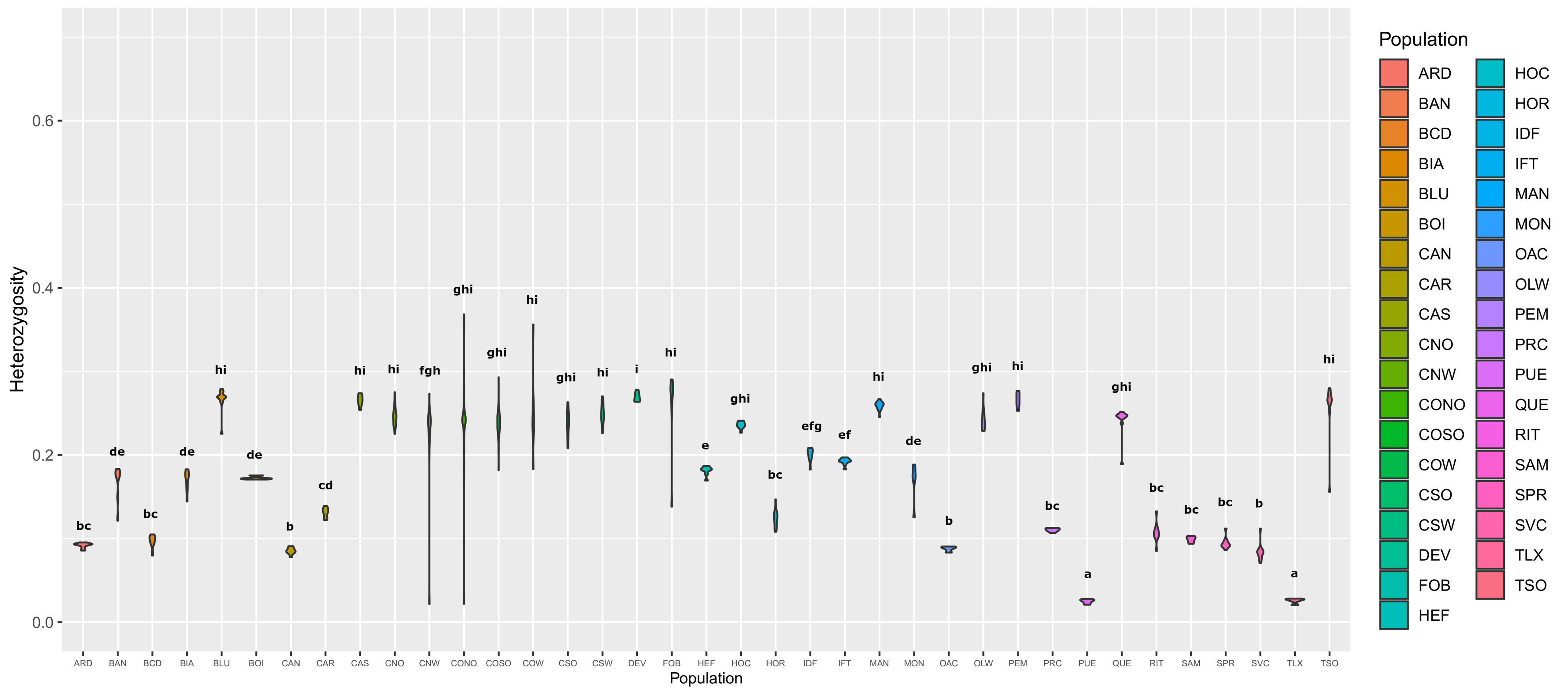

### Supplemental file 6

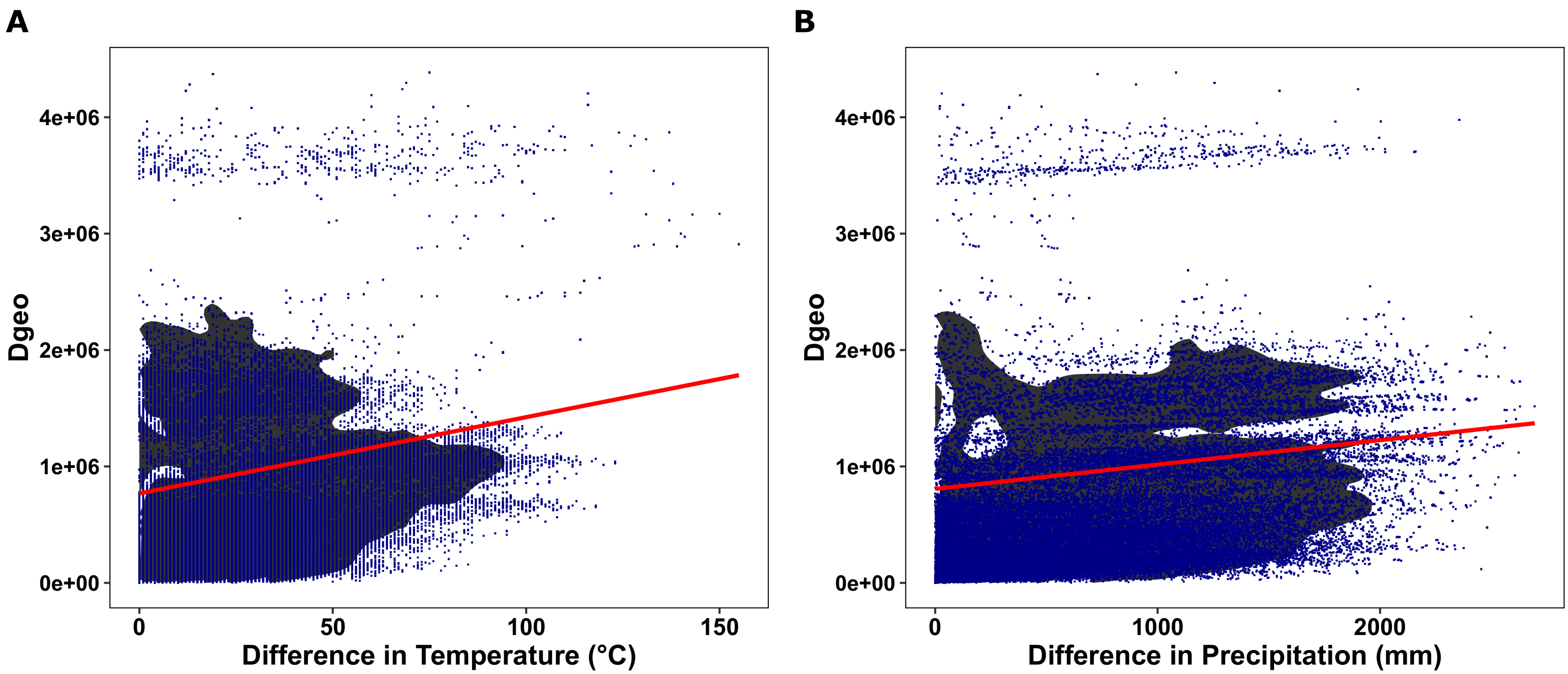

### Supplemental file 12

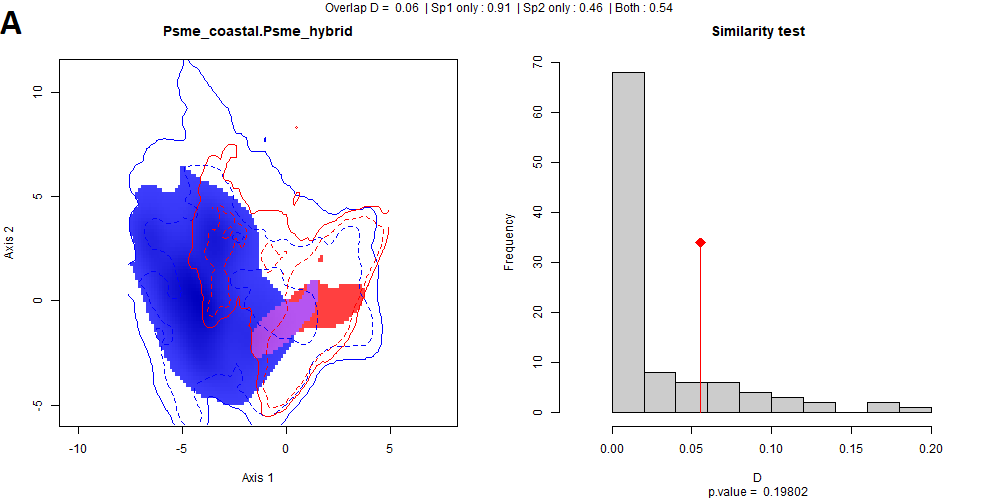

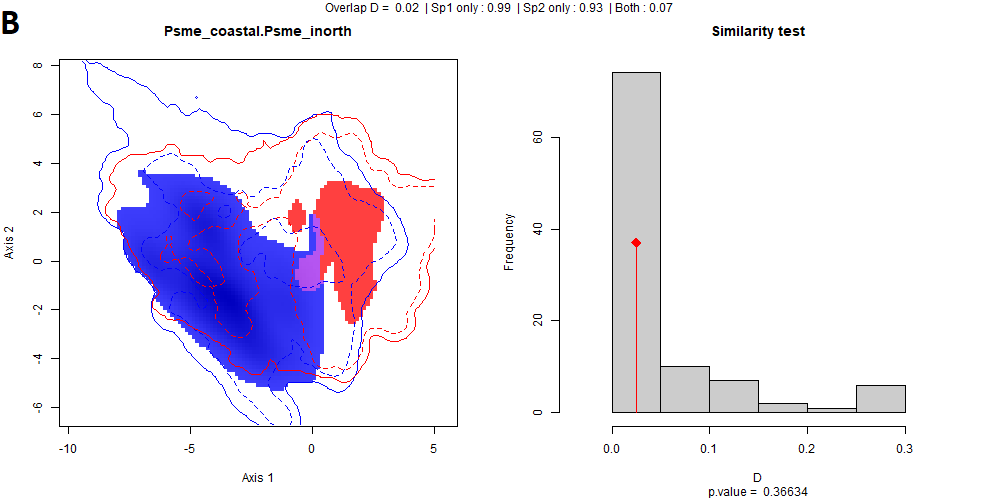

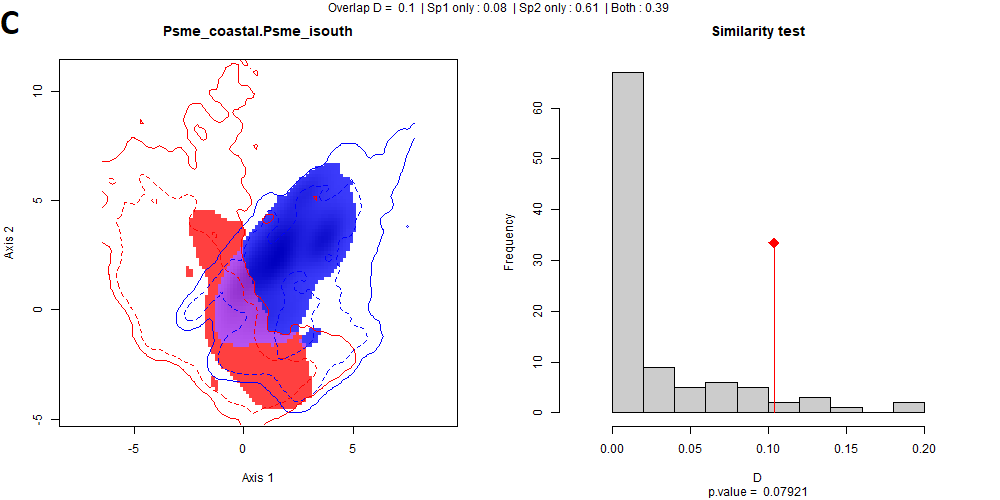

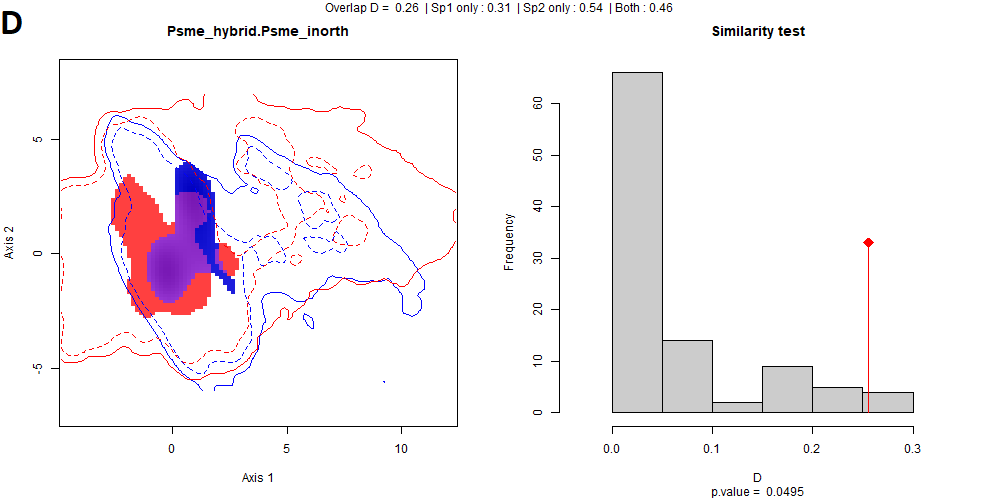

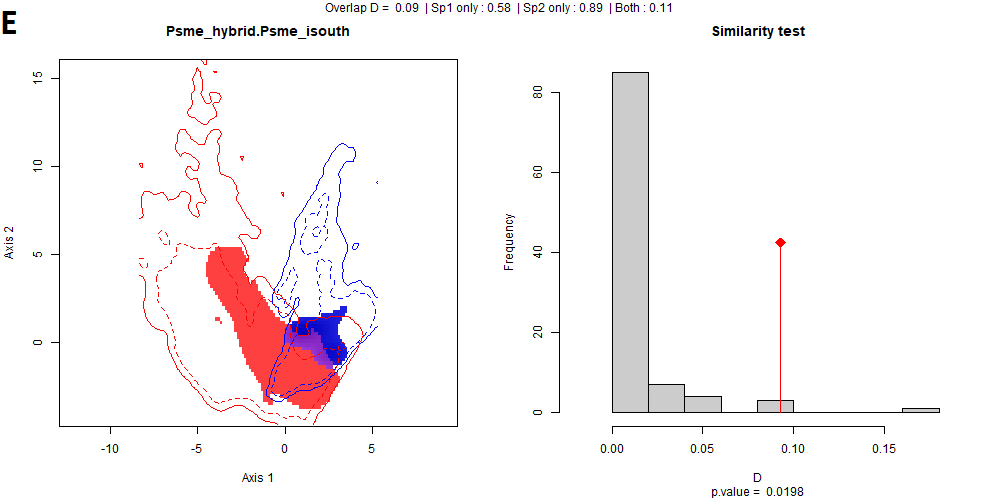

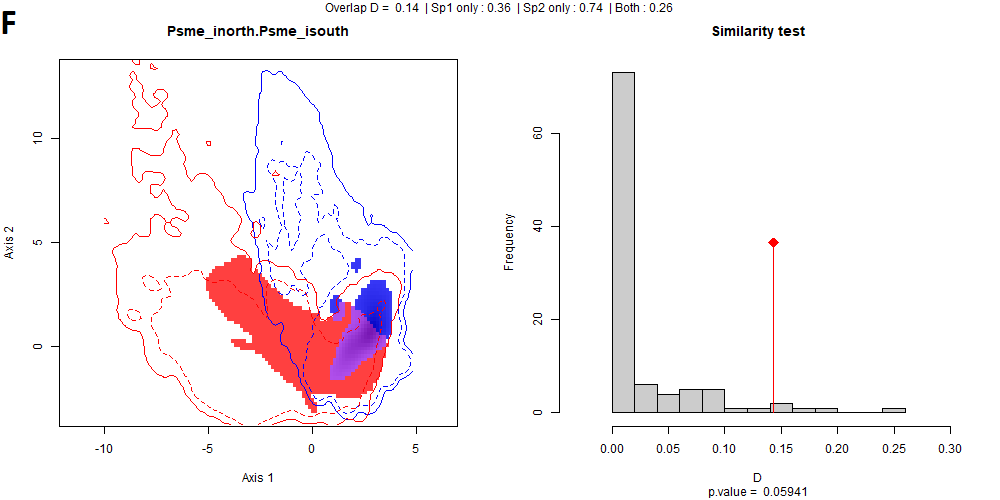

### Supplemental file 13

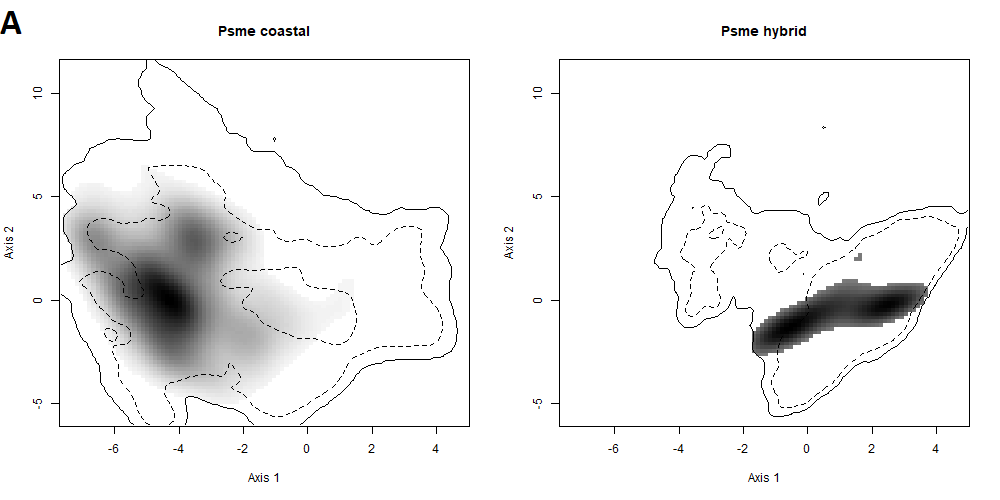

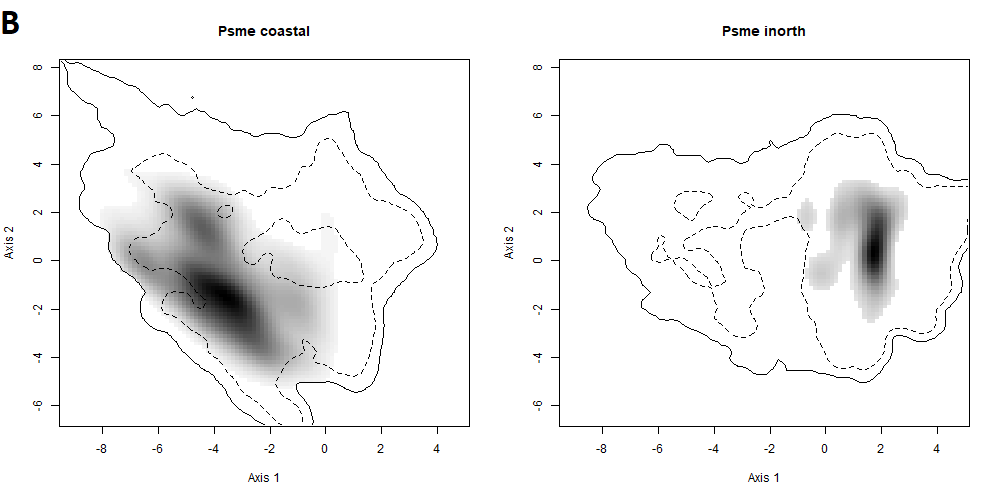

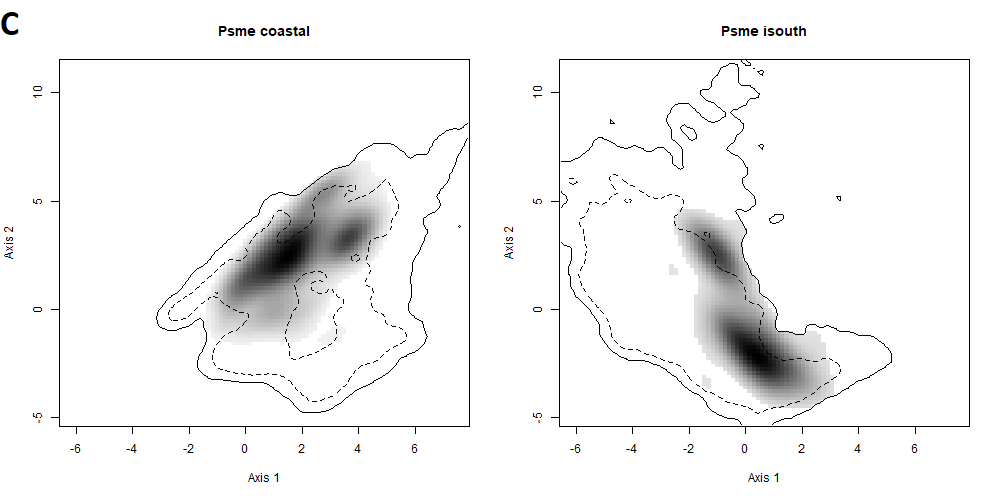

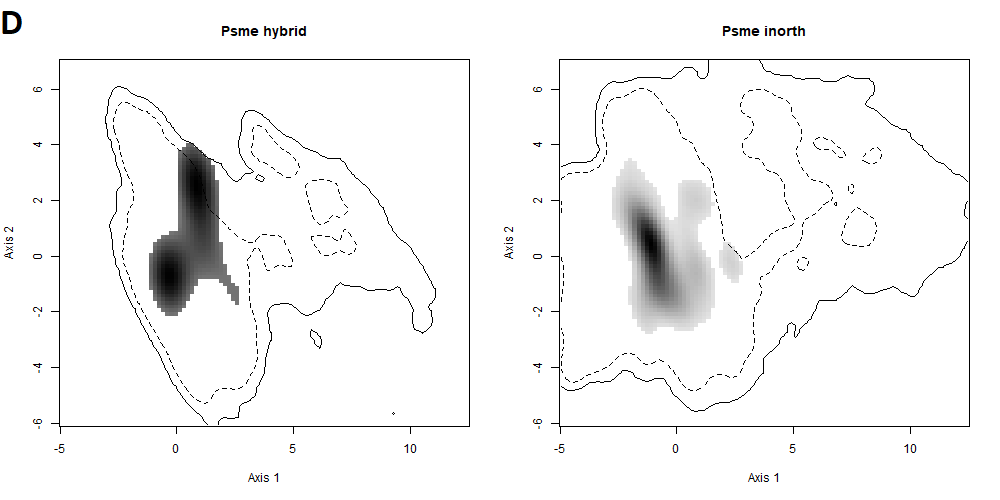

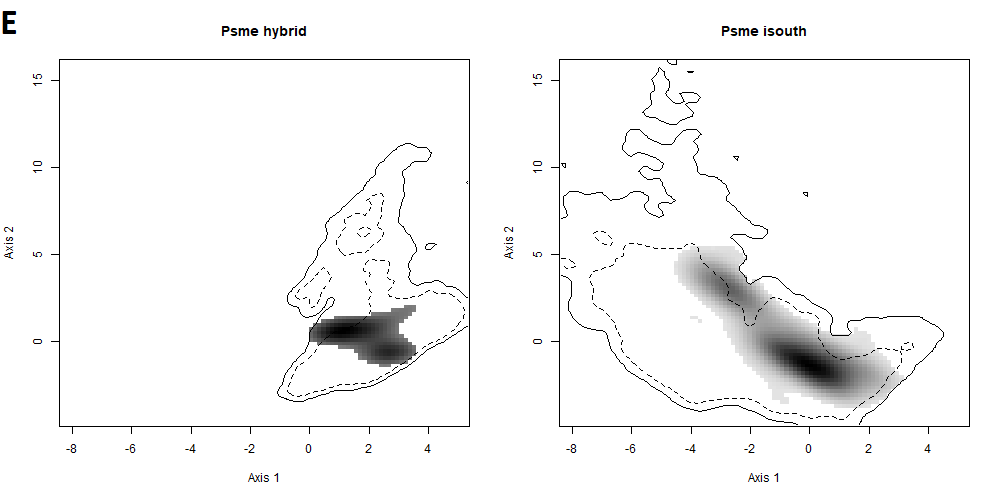

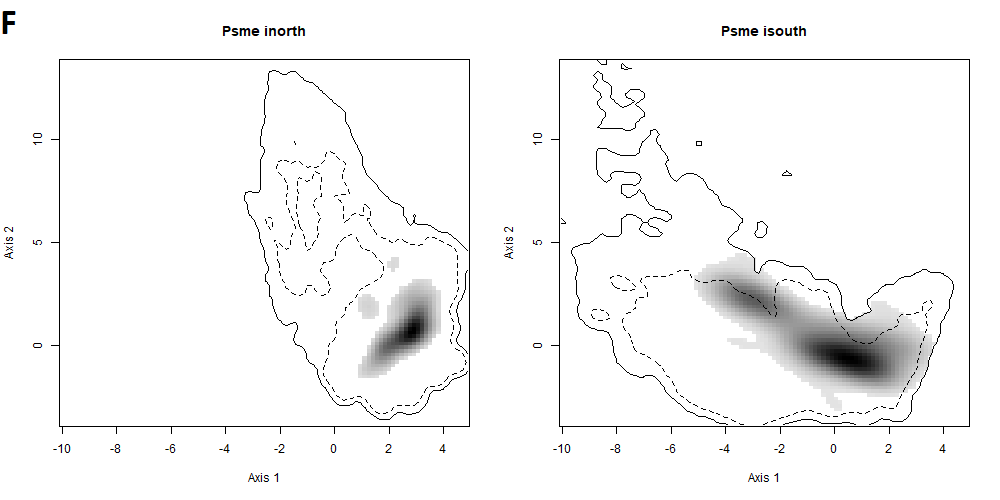
